# How human aging disrupts the head direction network: evidence from VR experiments and mechanistic models

**DOI:** 10.1101/2025.11.24.690256

**Authors:** Matthieu Bernard, Jonathan Shine, Andrej Bicanski, Thomas Wolbers

## Abstract

Navigational deficits during aging can severely limit mobility and reduce quality of life. While research on the underlying neural mechanisms has primarily focused on medial temporal lobe dysfunction, the head-direction (HD) system—a core component of the mammalian navigation circuit—remains largely unexplored in the context of aging. We established an immersive virtual reality paradigm that provides direct behavioral read-outs of HD signals. In addition, we developed a biologically inspired HD model, which accommodates noise sources that simulate age-related neural changes. Compared to younger adults, older participants exhibited larger angular errors, and a brief delay increased their heading uncertainty. In addition, our novel ring-attractor architecture shows that synaptic noise and small-scale neuronal loss replicate the magnitude and dynamics of the age-related deficits observed behaviorally. Together, these behavioral and computational findings provide the first evidence that aging compromises the fidelity and stability of the HD system. By pinpointing noise accumulation and neuron attrition as mechanistic contributors, our study significantly advances the understanding of spatial navigation deficits in old age, and it highlights novel targets for interventions aimed at preserving navigational abilities and quality of life.

## Introduction

Globally, the aging population is increasing at an unprecedented rate. By 2050, the number of people aged 60 and above is projected to reach over 2 billion^1^, and a significant proportion will have to cope with normative or pathological cognitive decline. However, one cognitive ability that is particularly relevant to everyday functioning has received relatively little attention in cognitive aging research: older adults can experience substantial declines in navigational abilities, for example, in finding their way in novel environments (for a review, see ^2^). Such deficits can severely restrict mobility and affect levels of (physical) activity and social participation. Moreover, reduced mobility and independence can increase a person’s risk of developing dementia, attesting to the importance of navigational ability for maintaining good health with increasing age^3^.

The ability to navigate one’s environment is underpinned by various neural populations and relies on different types of sensory information (for review, see ^4^). A central component of accurate wayfinding is knowing one’s orientation relative to the environment. In rodents, this knowledge is represented in a network of so-called head direction (HD) cells, which are defined as neurons that code for the animal’s facing direction when its head is oriented toward a specific direction relative to an external reference frame (e.g., North)^5–7^. More specifically, there is an increase in the firing rate of a HD cell when the head of the animal aligns with its preferred firing direction (PFD). HD cells have been found in the postsubiculum (PoS)^5^, the retrosplenial cortex (RSC)^8,9^, the anterodorsal thalamic nuclei (ADN)^10,11^, the entorhinal cortex (EC)^12^, as well as in the dorsal tegmental nuclei (DTN)^13^ and the lateral mammillary nuclei (LMN)^11^ in the rodent brain. Importantly, HD signals have also been identified in a multitude of animal models, including zebrafish^14^, fruit flies^15–17^, bats^18,19^, non-human primates^20^, and humans^21,22^.

To understand the mechanisms underlying HD cell properties, several ring attractor models have been proposed throughout the years^23–28^. These models replicated the properties found in the rodent (for review, see ^29^) to varying degrees. In contrast, others tried to link them to other spatial navigation features, such as cue combination^30,31^ or path integration^32^. In a study, Castagnero et al.^32^ modeled path integration using a generative model with young, older, and mildly cognitively impaired (MCI) participants. This paper is one of the few that tried to model the impact of aging and MCI, but no significant differences were found between young and older participants, except that MCI had different model parameters.

At present, however, it is unknown whether and how age-related neurodegenerative processes affect computations of the HD system. One way to model these processes is to increase synaptic noise and allow deviations from perfect connectivity within the model, which, in the extreme, could lead to individual neurons becoming essentially disconnected from the rest of the attractor circuitry (equivalent to deletions). In the first comprehensive modelling study of HD^24^, the effect of imperfect weights is touched upon. It was shown that initially, uniformly distributed HD converged to a few local minima over time, showing a reduction of representational space. However, no systematic investigation of the HD attractors under manipulations that reflect aging has been conducted thus far, or how this property translates behaviourally.

The canonical self-connected attractor can be interpreted as an elegant algorithmic solution according to Marr^33^. Assuming aging goes hand in hand with some neuronal loss, increased neural noise, and variations in synaptic connectivity (see below), it becomes essential to develop a biologically plausible model, approaching the implementation level in Marr’s terminology, since it is the implementation-level that should be affected by aging. The classical attractor model^24^ (also see ^23^) requires self-excitation with unreasonably precise connectivity that is extremely sensitive to perturbations. This has been partially addressed by architectures without recurrence^27,28^. Here, we show that a similar model architecture fares well under network changes we may deem approximations to the effects of aging.

Several physiological changes have been reported in older adults, such as synaptic changes^34^ or loss of cerebral volume^35–37^. However, this loss does not necessarily indicate neuronal loss, as many factors can contribute^38^. Nonetheless, neuronal loss might exist in some proportion^39^, in the vestibular nuclei^40–42^ or the thalamus^43,44^, two key structures of the HD network. In addition, studies have described a deficiency in the vestibular system with aging^45–47^, with evidence pointing to a loss of hair cells^48,49^, deterioration of vestibular pathways^50,51^, and reduced synaptic drive of projections from the vestibular system to the DTN^52^, which could produce a noisier input signal to the HD circuit.

Together, these factors could lead to defects within the HD network, and developing a new model could help understand their relative contribution. For example, age-dependent synapse changes can be modeled as persistent deviations from ideal connectivity or disconnections by changing the weights between neurons in the model. Removing neurons from the network could approximate age-related neuronal loss. Finally, the HD model could integrate vestibular deficits by increasing the noise level of vestibular inputs.

The present study aimed to investigate the effects of aging on HD coding and to identify mechanisms underlying potential deficits in older adults. To achieve these aims, we combined a behavioral study in immersive Virtual Reality (iVR) with a biologically plausible simulation of the HD system. Older and younger adults used iVR to learn and retrieve the position of a single distant landmark within a circular arena, either immediately or after a 20-second delay, by physically rotating toward the landmark (Fig. 1A-E). To preview, we found that (i) older adults performed worse than younger participants across rotation angles, and (ii) that the impact of a delay before responding was present in both populations. In the second part, we present a biologically plausible implementation of a HD attractor network, partly inspired by previous models^27,28,53^ (Fig. 1F-G). The architecture is based on a 3-ring model (Fig. 1F) where excitatory self-connections are removed, and instead, separate excitatory and inhibitory connections are used (conforming to Dale’s principle^54^). We characterize its operation and show that it can cope with weight noise, neuron deletion, and variations in synaptic activity, compared to the standard model of HD coding. We predict changes to HD tuning curves with age by changing the levels of each noise source. We also assess the reduction in representational space that could occur when the model is subjected to aging-like perturbations by modelling the behavioral experiment conducted in the first part. We found that the model could approximate results across young and old adults.

**Figure 1:**
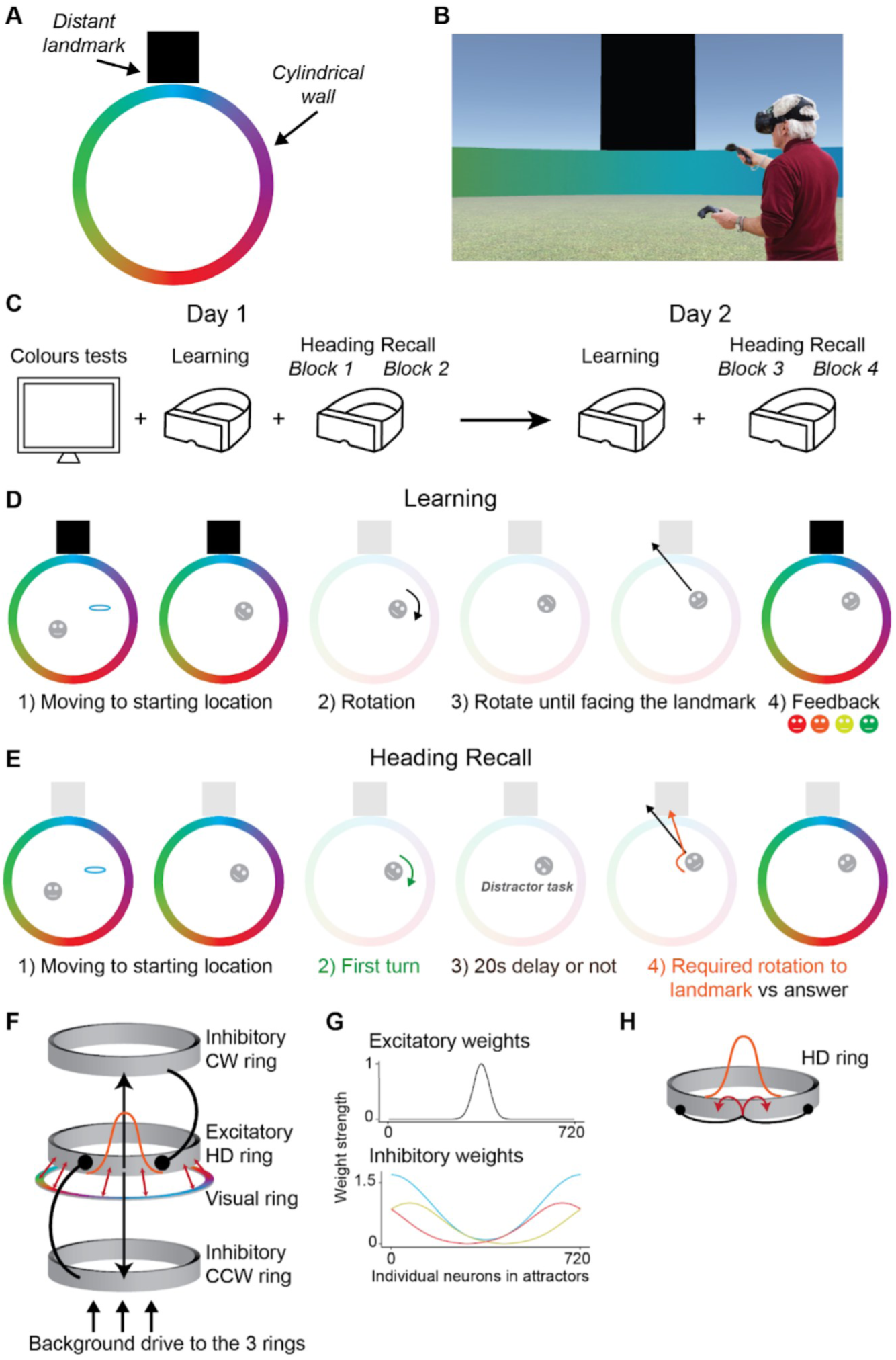
Virtual environment, behavioral task performed in immersive VR and model architecture. A, Schematic (birds-eye) view of the immersive virtual reality environment, consisting of a distant landmark and a cylindrical wall. B, Example view of a participant while performing the task in immersive virtual reality. C, Overall flow of the experiment across the two testing days. D, Design of the learning task with a trial example. E, Design of the Heading Recall task with a trial example. F, the 3-ring architecture with one ring exciting two separate rings that send back offset inhibitory connections (CW vs CCW weights). Modulating the connection strength of inhibitory weights translates the activity bump. Background drive is sent to all rings. A visual ring provides sensory feedback via one-to-one connections (red arrows) with the excitatory HD ring. G, excitatory (top) and inhibitory (bottom) weight distributions for a sample neuron in each ring. For inhibitory weights, gold corresponds to CW, red to CCW, and blue to a combination of both sets of weights. H, schematic of a single self-connected ring (red arrows) and inhibitory connections (symmetric and asymmetric) based on the HD model from Zhang (1996).

## Results

### Both age groups learned the landmark’s position

We first ensured that differences in colour perception did not confound our results and found no significant performance differences between age groups (see Supplementary Results). All participants in both age groups learned the landmark’s position (average error in the last five trials < 8°) on both days (Fig. 2A). An ANOVA on the number of learning trials revealed main effects of age (F(1,57)=12.39, p<0.001) and day (F(1,57)=9.79, p=0.003). In addition, older participants learned significantly faster on the second day than on the first (t(28)=-2.65, p=0.01). This was not the case for young participants (t(29)=-1.68, p=0.10). Note, however, that the ANOVA did not show an interaction between age group and day (F(1,57)=1.04, p=0.31), indicating that the change in learning speed across days did not differ between the groups.

**Figure 2:**
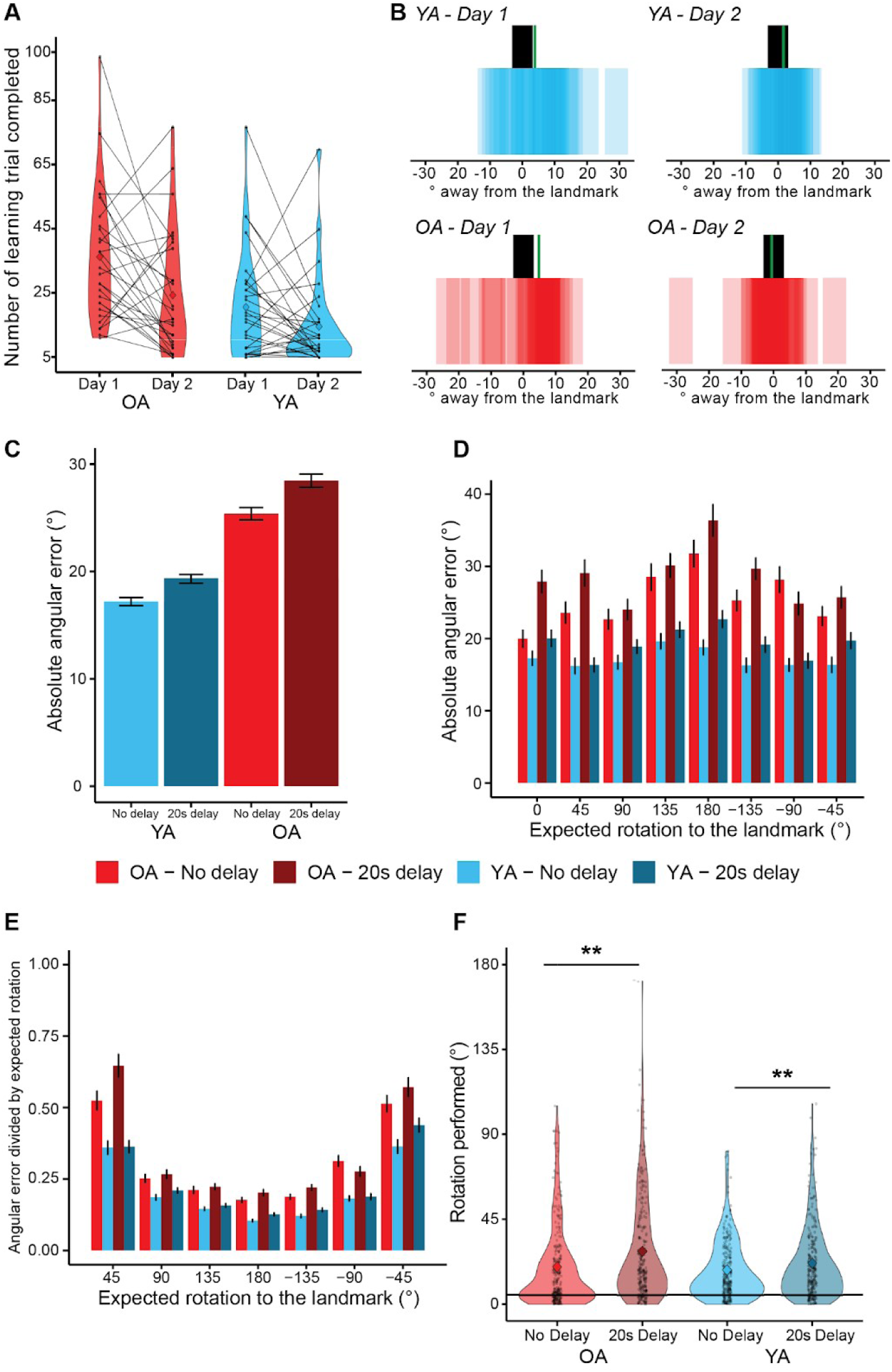
Behavioural results from young and older participants. A, Learning task: number of trials completed until passing the threshold for young (YA) and older (OA) adults on Day 1 and Day 2. Violin plots show the distribution of data. The dots represent individual data, and the diamond is the average across individuals. B, placement of the landmark by older adults (bottom) on Day 1 (left) and Day 2 (right). The black bar represents the correct position of the landmark, the coloured bars represent the individual data, and the green bar is the average answer from the participants. C-D, Heading Recall task: average absolute angular errors (coloured bars) for the two delay conditions. E, average ratio of the absolute angular errors divided by the associated expected rotation to the landmark for young and older adults for the two delay conditions.. F, average rotation performed between age groups and delay conditions. Violin plots show the distribution of data. The dots represent individual data, and the diamond is the average across individuals. The horizontal black line represents the threshold of ”Fixed heading” (i.e., 5°). In C-E, the vertical black lines represent the standard error of the mean. * p<0.05; **p<0.01.

In addition, we investigated whether there was a difference in landmark placement between age groups and across days (Fig. 2B). An ANOVA on landmark placement showed a main effect of days (F(1,57)=5.69, p=0.02), indicating that participants learned the landmark position faster on the second day. In contrast, there was no main effect for age group (F(1,57)=0.20, p=0.88) nor an interaction effect (F(1,57)=1.30, p=0.26). In general, both younger and older adults successfully placed the landmark within 20° of its real position on day 1 and 10° on day 2.

### Older adults show reduced heading accuracy and both groups are affected by delay

In the Heading Recall task, we did not find any performance differences across testing days (see Supplementary Results). Therefore, we pooled both days together for subsequent analysis. We first tested how age, delay, and expected rotation - defined as the angular turn required to face the landmark - affected heading accuracy in both groups. A three-way ANOVA on absolute angular error (Fig. 2D) revealed significant main effects for all three factors (age: F(1, 57)=9.83, p = 0.003), delay: F(1, 57)=36.93, p < 0.001), expected rotation: F(7, 399)=4.79, p < 0.001) and an interaction between delay and expected rotation (F(7,399)=3.21, p=0.003). For small expected rotations such as 0° or 45°, the delay had a more negative impact on the performance. In contrast, we found no interaction between age and delay (F(1,57)=1.12, p=0.29), age and expected rotation (F(7,399)=1.64, p=0.12), nor a three-way interaction (F(7,399)=1.87, p=0.072).

To evaluate whether performance was affected by the magnitude of the required turn, we expressed absolute angular error as a proportion of the corresponding expected rotation (Fig. 2E). Because this normalization is undefined for 0° trials, those data were analysed separately (see below). A three-way ANOVA on the resulting error ratio showed a main effect of delay (F(1,57)=24.34, p<0.001) and expected rotation (F(7,399)=297.51, p<0.003) but not age (F(1,57)=3.05, p = 0.086). Significant interactions emerged between age and expected rotation (F(7, 399) = 2.30, p = 0.002), and between delay and expected rotation (F(7, 399) = 22.38, p < 0.001), whereas the age × delay interaction did not reach significance (F(1, 57) = 3.64, p = 0.06). A significant three-way interaction among age, delay, and expected rotation was also observed, F(7, 399) = 3.50, p = 0.001.

Finally, we examined whether the magnitude of the initial turn, executed before the delay period, influenced performance (Supplementary Fig. 2). Larger first turns led to greater angular error, but only when the expected rotation was small (e.g., 45°). Once the expected rotation exceeded 90°, the size of the initial turn no longer affected accuracy. This pattern was comparable for young and older adults.

### Delay-induced performance decline in trials with correct initial orientation

In a subset of trials, participants already faced the landmark at the end of the first turn, hence the expected rotation was 0°. As a consequence, errors could be driven only by the initial turn or by the delay. The 20-second delay significantly broadened response variability (Supplementary Fig. 3) in both younger (t(29) = 2.83, p = 0.008) and older adults (t(28) = 3.09, p = 0.004). Quantifying rotation across delay conditions (Fig. 2F) revealed a significant main effect of delay (F(1, 57) = 14.82, p < 0.001), with both groups making larger turns (YA: t(29) = –3.12, p = 0.004; OA: t(28) = –2.29, p = 0.03) after the delay, but no main effect of age (F(1, 57) = 0.0008, p = 0.98) or interaction (F(1, 57) = 0.91, p = 0.34). For further results, see Supplementary Results.

### Delay period increased error variability without inducing response clustering

We reasoned that age-related noise might collapse the head-direction attractor into a few preferred headings, which would appear behaviourally as tighter clustering (lower circular SD) after a delay. Therefore, we quantified response clustering with the circular standard deviation (CSD; Supplementary Fig. 6A). A three-way ANOVA on CSD revealed significant main effects of age (F(1, 57)=16.93, p < 0.001), delay (F(1, 57)=11.52, p = 0.001), and expected rotation (F(7, 399)=2.56, p = 0.014). A delay × expected-rotation interaction was also significant (F(7, 399)=2.62, p = 0.012), whereas the age × delay (F(1,57)=0.12; p=0.73) and age × expected-rotation (F(7,399)=0.52; p=0.82) interactions were not. Contrary to our “reduced representational space” hypothesis (and classical HD models), the CSD increased—rather than decreased—after the 20-s delay, indicating that responses became more dispersed. The same pattern emerged when trials were stratified by the size of the initial turn (Supplementary Fig. 6B-D), further arguing against a delay-induced clustering of headings in older adults.

In summary, we found that on average, older participants performed worse than younger participants. In addition, the delay period negatively affected both age groups, and the expected rotation amount affected performance. Specifically, an accumulation of noise was present during the rotation of the answer phase. However, when the heading of the participant was fixed before answering, older participants performed worse after the delay period.

### The 3-ring model is robust to aging-like changes

To gain a deeper understanding of the neural mechanisms that could explain our behavioral findings, our modelling analysis proceeded in three steps. First, we quantified how each noise source affects HD stability in static simulations. Second, we assessed how these perturbations influence the HD ring during the trial phases of the behavioural task. Third, we evaluated combinations of noise sources to determine which aging-related perturbations best reproduce the human behavioural data.

We implemented the 3-ring model of the HD circuit (Fig. 1F and Methods), with one excitatory ring projecting to two inhibitory rings, which project back (CW and CCW, respectively) to the excitatory ring. Next, we characterised it by 1) showing that the model could accommodate neuron deletion while the recurrent HD attractor could not (Fig. 3A); 2) testing whether the HD bump could follow participants’ trajectories (Fig. 3B). For more details, see Supplementary Methods.

**Figure 3:**
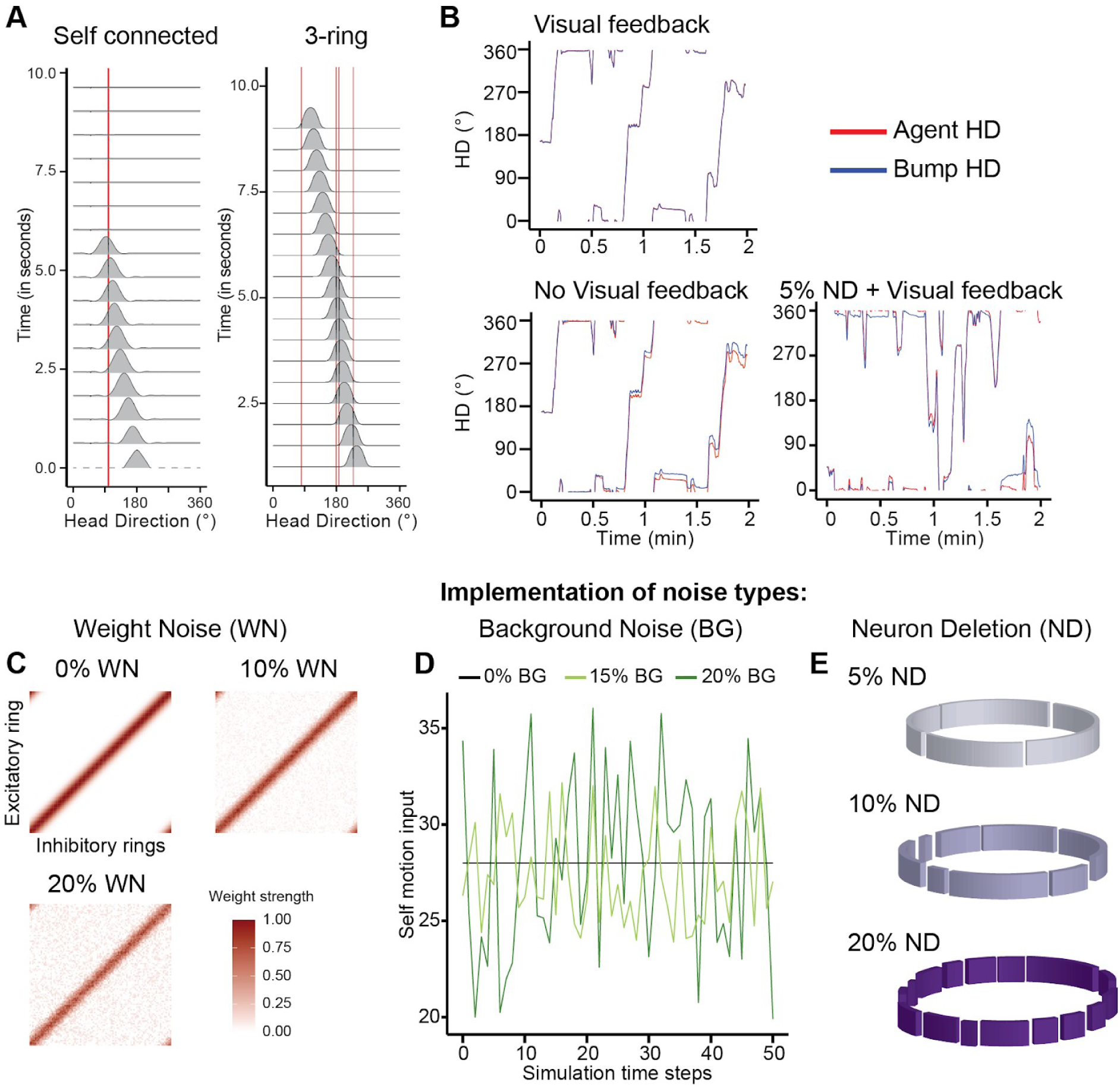
Architecture and characterisation of the model. A, left: simulation of self-connected attractor model after removing a single neuron (red line) and letting the bump move in one direction. Upon reaching the dead neuron, the bump collapsed. On the right, simulation of the model developed in this chapter after adding 1% neuron deletion (red line, indicating dead neurons in the HD spectrum). The bump passed all dead neurons and maintained its shape. B, Trajectories performed by the agent when visual feedback was present (top right) or not (bottom left) and when 5% of neuron deletion was introduced (bottom right). C, weight matrices as heatmaps for 0, 10, and 20% weight noise (WN). D, example of background drive values to the rings (i.e., self motion input) for 50 stimulation time steps for 0, 15, and 30% background noise. E, schematic of the ring attractor after applying neuron deletion.

Next, we ran simulations to examine the influence of individual noise sources (Fig. 3C-E) on the HD bump shape at rest (i.e., no velocity input) for 2 minutes. For weight noise (Fig. 3C), we found a fuzzier bump, and HD slowly drifted to another position after a few seconds (Fig. 4A). There was no consistent drift as a function of background drive (Fig. 3D), but we observed jitter in the HD and an increase in bump width (Fig. 4B). For deletions (Fig. 3E), the bump widened (from 47° at 5% to 58° at 20%), and drift was observed, similar to weight noise (Fig. 4C).

**Figure 4:**
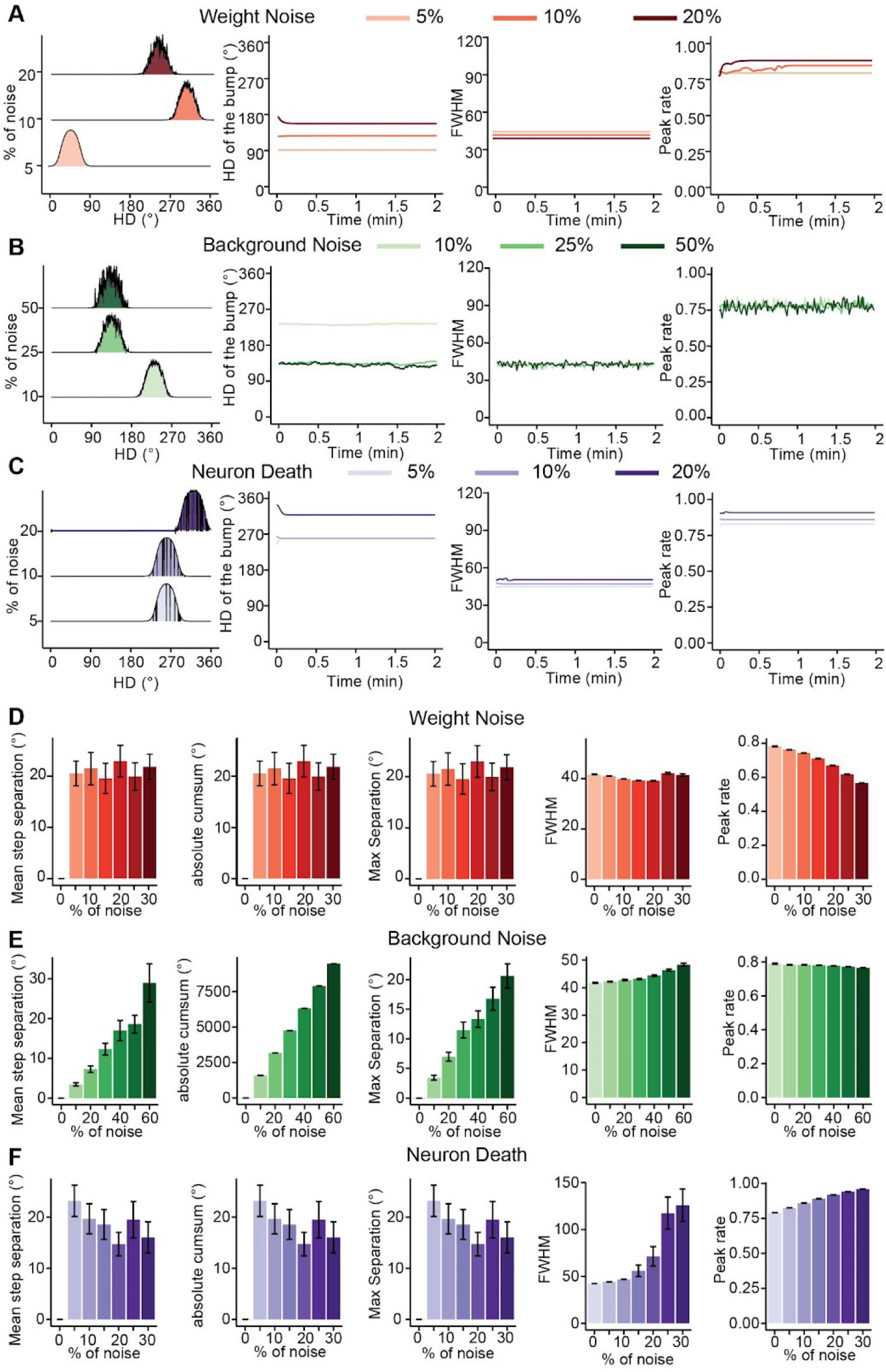
Influence of noise levels on the tuning curve shape and behaviour of the HD model. Measures of the tuning curve for (A) noise on the weights, (B) noise in the background drive, and (C) neuron deletion. Left to right, tuning curves from the excitatory ring for different noise levels after 1 minute of simulation; the HD over time; the FWHM over time; and the peak rate over time. Measures of the behaviour of the HD model for (D) noise on the weights, (E) noise in the background drive, and (F) neuron deletion. Left to right, the average step separation over simulation steps; the absolute cumulative sum of HD changes; the maximum separation from the initial HD and the final HD; the average FWHM; and the average peak rate. Simulations lasted 2 minutes, and no velocity input was given. The black lines represent the standard error.

To characterise those changes in more detail, we ran 50 simulations with a broader range of noise levels. Each simulation lasted two minutes and did not include velocity input. For weight noise (static, synaptic weight perturbations), the cumulative sum of changes indicated a random drift to a stable position (Fig. 4D). With increasing weight noise, the FWHM remained stable, and peak rate decreased (Fig. 4D). The average magnitude of drift (maximum separation) remained approximately constant (Fig. 4D).

For increased variability in the background drive, larger amplitudes in the mean step separation, absolute cumulative sum, and maximum separation at higher noise levels were observed (Fig. 4E). In addition, the absolute cumulative sum was greater than the step cumulative sum (Fig. 4E). This is due to the shape of the bump being fuzzier, which modifies the heading at every step when calculating the cumulative sum (as seen in Fig. 4E). The FWHM increased across noise levels while the peak rate decreased.

Finally, as we increased the percentage of neuron deletions, we did not observe a specific pattern of change in HDs (Fig. 4F). This result was due to the random drift to the stable position at the start of the simulation. However, we noticed a significant increase in FWHM at higher levels of neuron deletion, indicating that, while HD could still be adequately represented under these conditions (provided feedback anchors the signal to the external world, cf. Fig. 3B), the precision of the heading representation is adversely affected (Fig. 4F). We also noticed an increase in the peak rate for high noise levels.

In summary, each type of noise influences the HD network in a specific manner. Weight noise induced a fuzzier bump with gradual drift and widening, accompanied by reduced peak rate. Background drive noise led to similar bump widening, with increased drift and fuzziness, as well as a decrease in peak rate. Neuron deletions caused random drift to a stable position, with higher deletion levels significantly increasing bump width and reducing precision, though the HD representation remained intact. Across tested noise levels for each type, the HD bump remained present, indicating that the 3-ring model is robust to age-related changes and can be used to investigate them. Most importantly, such a highly perturbed HD system could still convey reliable HD signals when supplied with visual feedback (Fig. 3B).

### Noise types differentially affect simulated task performance

Next, we implemented the behavioral task (i.e., Heading Recall) in the model and examined how each noise type affected the agent’s performance. Individual trials were timestamped according to the trial phases (see Methods section and Fig. 5A). Each trial began with visual feedback aligning the HD bump to the agent’s heading (Phase a). The agent rotated to the start orientation (Phase b), paused (Phase c), turned with visual input turned off (Phase d), underwent a 0- or 20-s delay (Phase e), and finally rotated to face the landmark, aligning the HD ring with the landmark’s color (Phase f). To assess the impact of the noise, we calculated the angular error (the difference between the agent HD and the bump HD at each timestep) and the average rate of change during each phase (derived from the difference in angular velocity between the agent and the bump).

**Figure 5:**
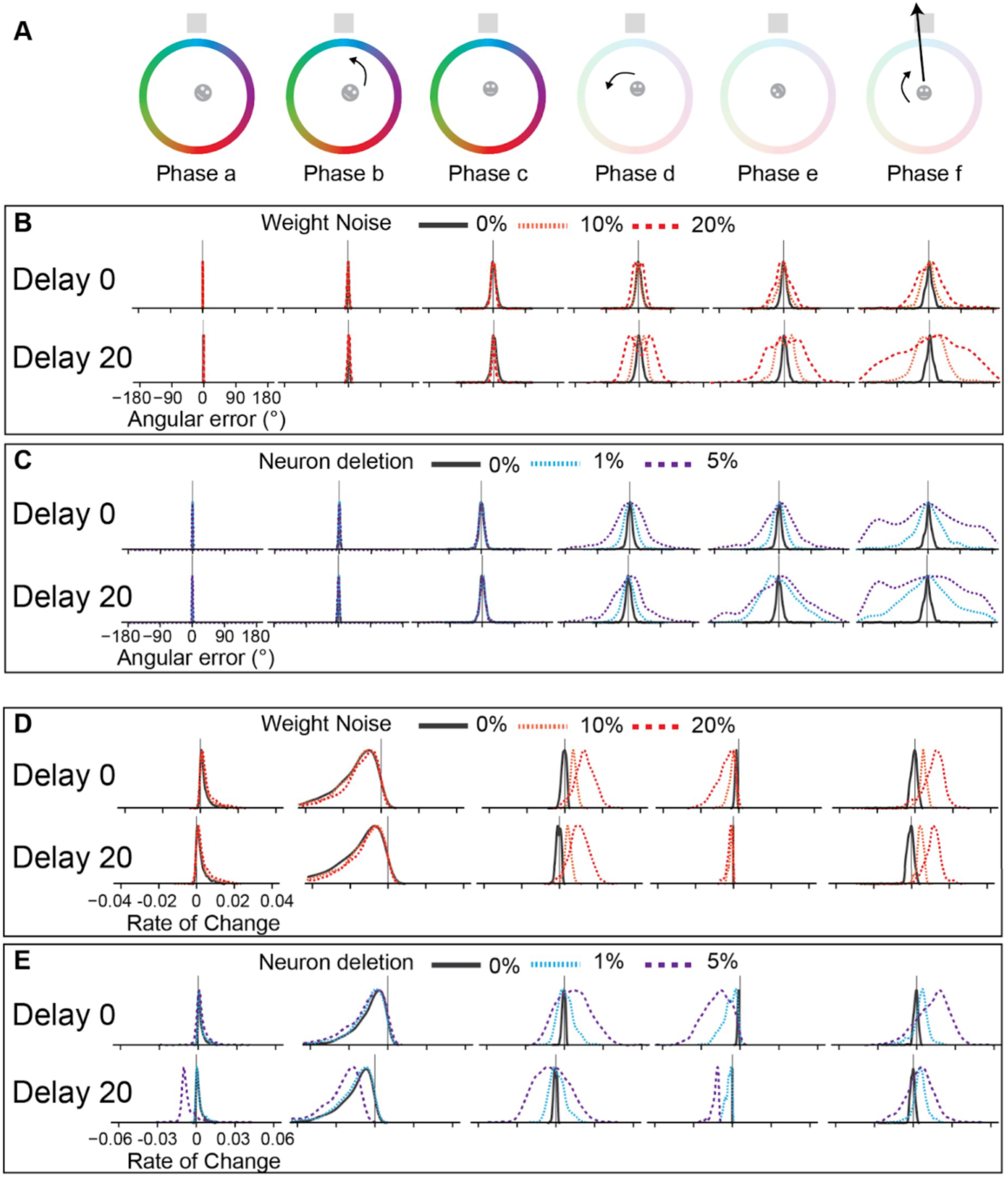
Evolution of angular error and rate of change for increased weight noise and neuron deletion. A, the different phases forming a single trial executed by the agent. Phase a, reset of the network; Phase b, rotation toward starting orientation; Phase c, facing starting orientation; Phase d, performing the first turn; Phase e; delay condition; Phase f; answer from the agent. B,C, the density plots show the distributions of the angular error (difference between agent HD and the **HD** indicated by the network at the end of a phase) measured at the end of the trial phases for weight noise (B) and neuron deletion (C). The vertical black line represents 0. The plots were separated by level of noise (colours), by phases (from left to right), by delay conditions (top - no delay, bottom - 20s delay). D, E, the density plots show the distributions of the rate of change (average of the difference in velocity of the **HD** of the agent and the bump) measured for weight noise (D) and neuron deletion (E). The vertical black line represents 0. The plots were separated by level of noise (colours), by phases (from left to right), by delay conditions (top - no delay, bottom - 20s delay).

### Weight Noise

First, we compared the angular error in the simulated behavioral task for different levels of weight noise (static perturbations of synaptic strength). Resetting of HD worked over trials, excluding a potential accumulation of error within the system across trials, as shown by the HD peaks centered on zero angular error in Fig. 5B (Phase a). We found increased angular error after the first turn (Phase d, Fig. 5B) and the answer phase (Phase f, Fig. 5B) for both delay conditions. In addition, the 20-second delay condition produced a substantial error (Phase e, Fig. 5B) not present in the no-delay condition. This was in part due to drift during the delay phase, which is exacerbated by higher levels of weight noise (compare Phase e with and without delay in Fig. 5B). This is confirmed by looking at the rate of change during the delay phase (Phase e, Fig. 5D). Indeed, there was a higher rate for larger noise, indicating that the HD bump was drifting since the real HD was constant during this phase.

Furthermore, we noticed an overall increase in the rate of change without visual feedback (Phases d and f) for higher noise levels (Fig. 5D). This increase was positive, meaning that the agent’s velocity was larger than the translation speed of the activity bump for higher level weight noise.

### Background drive

We did not find a systematic increase in angular error with higher background drive noise levels in either delay condition (Supplementary Fig. 7). The spread of angular errors was due to the loss of visual feedback from Phase a to Phase c (Supplementary Fig. 7B). However, we found a negative rate of change across the phases, with its fastest during the delay phase (Phase e, Supplementary Fig. 7C). This rate of change was due to the network’s erratic bump, which had an unstable HD compared to the agent’s heading. Indeed, our static simulations showed that the HD of the bump varied over time, while the average HD remained at the same position. Those changes in HD would create faster velocity values than the agent’s, producing the negative rate of change values in Supplementary Fig. 7C.

### Neuron deletions

Finally, we assessed the impact of deletions on the performance of the simulated behavioural task. We found a large spread of angular errors at every phase without visual feedback (Phases d, e, and f, Fig. 5C). Based on our previous static simulations, we conclude that the spread resulted from a higher probability of getting stuck in local minima during the trials. Analysing the rate of change across the different phases confirmed that the HD bump can get stuck in local minima. During the delay phase (Phase e, Fig. 5E), we observed a constant negative rate of change, with the HD bump moving toward the stable position, similar to the weight noise. Interestingly, the rate of change was faster and had a larger spread during the no-delay than during the 20-second delay period. Since the measure was an average over the entire phase, this indicates that the bump did not reach the stable position during the no-delay phase and continued to move. During the 20-second condition, the bump had enough time to reach a fixed position and stay static, decreasing the average rate of change.

Further evidence from the answer phase (Phase f) showed that most trials had a positive rate of change, especially for the 20-second delay (Fig. 5E). These results imply that the HD bump frequently settled into local minima during the delay phase. As a consequence, its speed during the response phase was reduced because it had to overcome and escape these attractor basins before matching the agent’s velocity. Lastly, the first turn phase (Phase d) distribution was dispersed on both sides, with positive and negative rate-of-change values (Fig. 5E). This range indicated that the bump was not stuck in local minima in every trial.

### Adding neural deletions to other sources of noise provides a better fit for older adults

After investigating each noise type individually, we combined the noise types (Fig. 6) to identify the set of perturbations that best accounted for the behavioural differences between age groups. Model–data correspondence was assessed using two complementary approaches (see Supplementary Methods for details): (i) comparison of descriptive statistics and (ii) log-likelihood estimates obtained via kernel density estimation. Both approaches yielded broadly convergent, though not identical, patterns.

**Figure 6:**
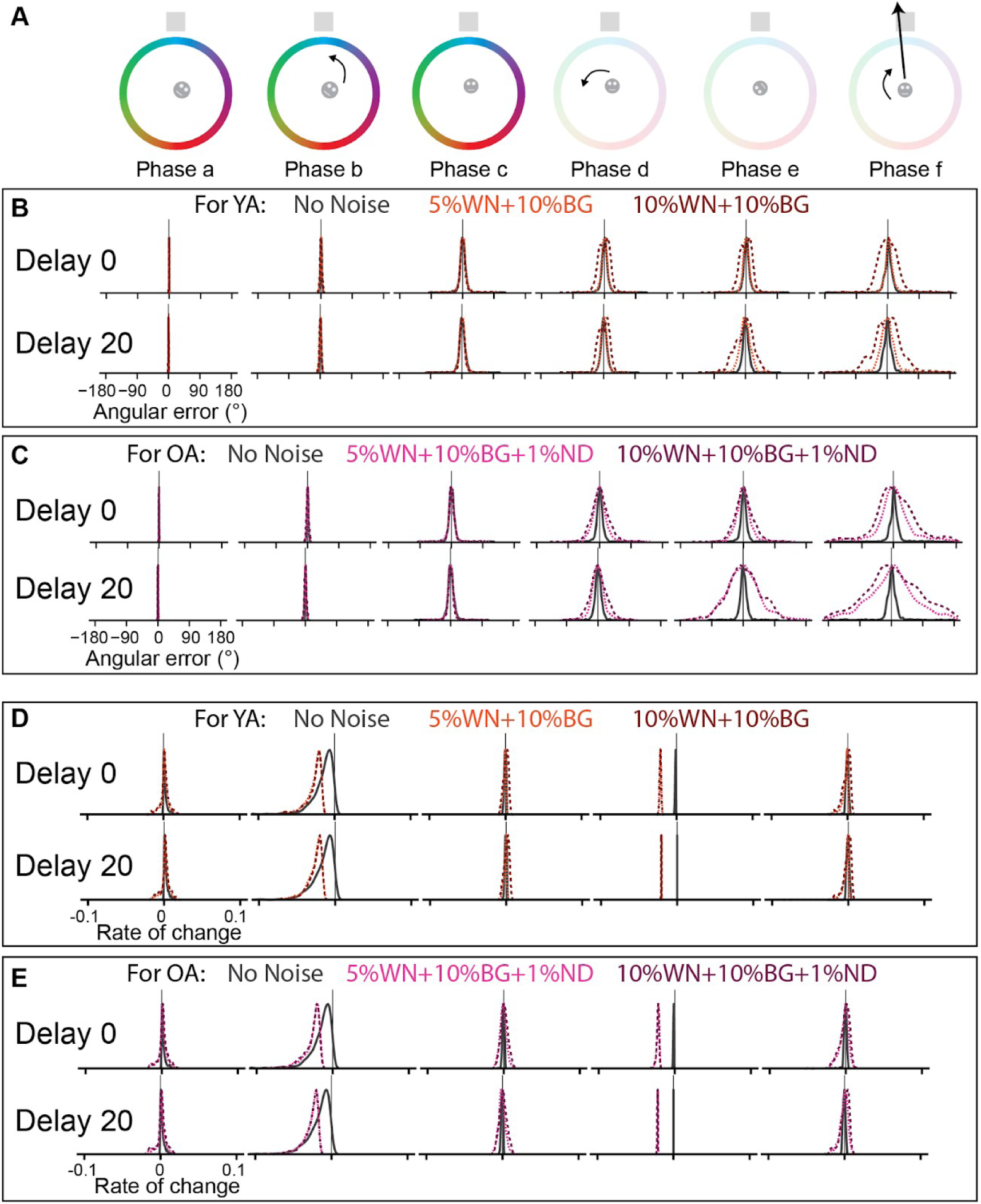
Evolution of the angular error and rate of change for noise combination testing behavioural data. A, the different phases forming a single trial executed by the agent. Phase a, reset of the network; Phase b, rotation toward starting orientation; Phase c, facing starting orientation; Phase d, performing the first turn; Phase e; delay condition; Phase f; answer from the agent. B,C, the density plots show the distributions of the angular error (difference between agent HD and the HD indicated by the network at the end of a phase) measured at the end of the trial phases for noise combination testing young adults (B) and older adults (C). The vertical black line represents 0. The plots were separated by level of noise (colours), by phases (from left to right), by delay conditions (top - no delay, bottom - 20s delay). D, E, the density plots show the distributions of the rate of change (average of the difference in velocity of the HD of the agent and the bump) measured for noise combination testing young adults (D) and older adults (E). The vertical black line represents 0. The plots were separated by level of noise (colours), by phases (from left to right), by delay conditions (top - no delay, bottom - 20s delay).

Descriptive statistics indicated similar levels of background drive variation and synaptic perturbations in both age groups, but older adults were better characterised by models that additionally incorporated mild neuron loss. Consistent with this, the kernel density estimates showed that models incorporating small amounts of neuron deletion produced higher log-likelihoods for the older-adult condition, indicating a better fit (Supplementary Table 5). Notably, the best-fitting models for older adults required less synaptic noise than those for young adults (Fig. 7) - a counterintuitive result suggesting that neuronal loss itself may introduce sufficient instability, reducing the need for additional synaptic perturbations to explain the behavioural variance. See Supplementary Results for more details.

**Figure 7:**
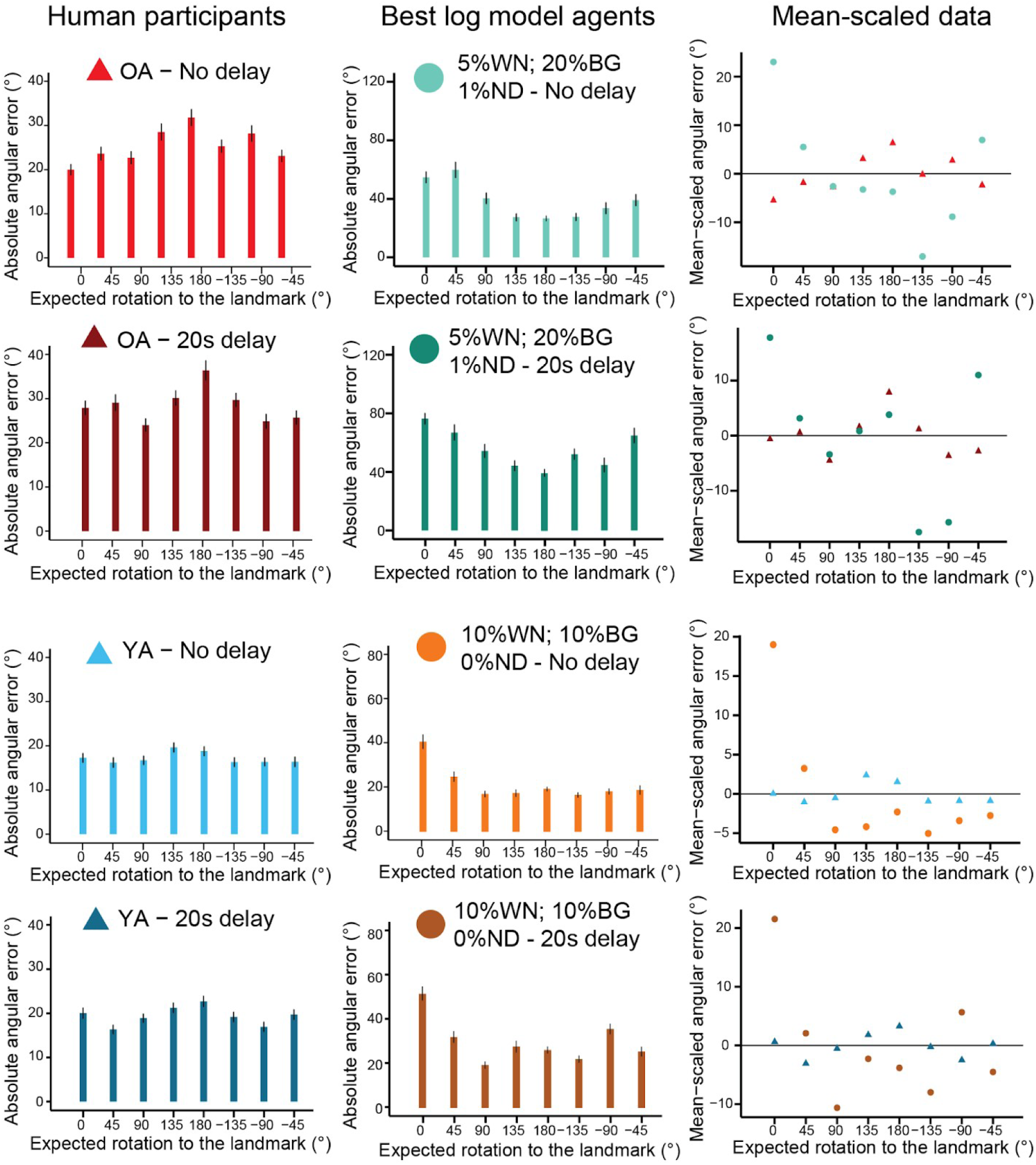
Performance comparisons between human participants and model agents with the best log-likelihood results. Left, Average absolute angular errors (coloured bars) for young (YA) and older (OA) adults for the two delay conditions. Middle, Average absolute angular errors from the combined noise-level configurations that return the best log likelihood-results in the kernel density analysis (corresponding to the final angular errors in Phase F of Fig. 6). The vertical black lines represent the standard error for each bar. Right, Mean-scaled angular errors corresponding to the bar plots shown on the left and middle.

## Discussion

Understanding how aging affects the neural computations supporting navigation remains a central challenge for cognitive neuroscience. Here, we combined an immersive VR experiment with a biologically grounded HD model to identify which aging-related neural perturbations best explain behavioural deficits. Participants first learned the angular position of a distant landmark until meeting a predefined performance criterion. At test, they were rotated to different starting orientations and then had to turn until they faced their remembered position. As predicted, older adults showed reduced heading accuracy and increased drift, particularly following a delay. To gain a deeper understanding of the deficits in the underlying neural computations, we complemented the experiments with a HD model based on a 3-ring architecture, driven by tonic vestibular input. Specifically, this allowed us 1) to explore three distinct noise sources that may compromise the aging HD system, and 2) to contrast our 3-ring model with the canonical self-connected ring attractor. By combining different noise sources and comparing the modelling results with human data, we found that weight noise and background drive were present in young adults, while the addition of minor neuron deletion best explained older adults’ performance. Interestingly, both behavioral and modelling results did not support the idea of reduced representational space in HD. Rather, the multi-ring model suggests that local minima are counteracted by variable, tonic vestibular drive, a direct consequence of adopting a biologically plausible architecture that necessitates tonic vestibular inputs^55,56^, compared to implausibly precise recurrent networks. The latter also violate Dale’s principle, do not naturally accommodate vestibular drive, and require precisely tuned connections.

At the behavioural level, the pattern of angular errors was proportionally higher for smaller expected rotations to the landmark (45°) than for large ones (180°). In addition, older participants generally performed worse than younger participants, making larger angular errors overall. In addition, the 20-second delay negatively impacted both age groups, who also showed a wider range of answers after the delay period. Thus, contrary to predictions from classical self-connected attractor models, aging did not collapse the HD representation into fewer stable orientations after the delay. Instead, we observed increased drift rather than clustering. Importantly, our results are unlikely due to a deficit in learning, as both age groups performed extensive learning and passed a strict performance threshold of 8°, and we did not find any age difference when participants placed the landmark back after a block of trials.

Separating the participants’ answers by the expected rotation to the landmark suggested that higher angular errors occur with larger rotations (e.g., 180°), especially in older adults. However, this measure did not account for the variable amount to rotate, as participants faced different orientations before answering. When computing a comparable metric, we observed that greater errors would come from a smaller expected rotation (i.e., 45°). Taken together, those results suggest that the main source of errors for larger expected rotation (from 90° to 180°) is the accumulation of noise during the execution of the answers. For the smaller expected rotation trials (45°), the error might be due to increased noise during the completion of the first turn. In addition, for 90° to 180° turns, the magnitude of those turns was greater. Then, the noise from the velocity integration could accumulate longer than the 45° expected rotations during the answer phase. Previous studies on motion perception in humans found similar results where participants had a systematic bias in their velocity integration during passive translation^57,58^. For active translation, path integration tasks also showed similar results, where noise accumulation played a significant role in the error^59^.

To determine which aging-related neural changes could produce these behavioural signatures, we established a biologically grounded HD model. The seminal model by Zhang^24^ only touched upon the influence of some noise on the weights. However, their elegant algorithmic solution required revision when assessed at the level at which aging manifests, i.e., the implementation level. Specifically, the canonical self-connected network is not designed to accommodate certain types of noise, such as neuron deletion, because the HD bump can collapse with as little as one dead neuron. To avoid this problem, we adopted a 3-ring architecture, supported by anatomical evidence^60^ concerning HD cells in the DTN and LMN. Indeed, lesions to the PoS^61^ or EC^62^ do not disrupt the HD signal in ADN or LMN. In contrast, lesions of the semicircular canals^63,64^ or bilateral LMN^65–67^ prevent the generation of the HD signal in the ADN or PoS. Those results suggest that the HD signal depends on the vestibular system and is generated in the LMN, then projected to the ADN before reaching cortical regions. This architecture removes the need for direct recurrence and enables the study of neuron deletion, strong perturbations to synaptic weights, and the incorporation of vestibular drive. Most notably, with suitable feedback, even a damaged attractor can still function. Clustering (reduction of representational space) was counteracted by variations in the vestibular background drive, allowing the HD to escape attractor basins, consistent with the experimental data. Note that this follows naturally from the inclusion of background drive, which - in our model - is needed for a bump to form in the first place, and renders the model consistent with the HD literature^63,64^. This result also suggests that older adults’ HD systems may be able to handle increased instability from the background drive arising from the vestibular regions. Alternatively, it could be framed as a compensatory strategy to avoid getting stuck in local minima. We also suggest that increased neuron deletion will manifest in less precise HD coding, here quantified as higher FWHM and peak rate in individual HD tuning curves.

The decision to add different noise sources was motivated by the current state of the literature on the impact of aging. Each noise type was intended to capture a specific physiological change known to affect the aging brain. Weight noise modelled synaptic change, where connections between neurons would be impaired^34^. Neuron deletion was linked to cerebral volume loss in brain regions that are presumed to contain HD cells^35,36,43^. Background drive was associated with vestibular deficiency, as both are the generating power of the HD system^45–47^. To the best of our knowledge, our model is the first to address this question directly. However, the exact connectivity and architecture of the human HD system remain unknown. Thus, our exploratory model may require revisions in the future. Specifically, we could not replicate the full range of errors we saw within our older group. Similarly, we expected the simulated aging group to show a better fit to the data at a higher level of synaptic noise compared to the simulated young group. Additional brain mechanisms could reduce the impact of cerebral loss but were not modelled^68,69^. An extended model could compensate for the loss of neurons through some form of homeostatic regulation and plasticity, or recalibration^70^, which provides a good starting point for future work.

In the context of aging, it is interesting to qualify statements about the present model by comparing it to the well-characterized HD system in insects (e.g., the Drosophila circuit^15–17^). We hypothesize that it should be more susceptible to neuron deletion because robustness to perturbations due to aging is not a concern in insects with shorter lifespans. This is reflected in a lower number of neurons compared to mammals like humans and rodents, which have a longer life span, suggesting neural circuits must be able to accommodate more neuron deletion and perturbations of synaptic connectivity^34^. This is consistent with the effect of network size we observe in the present model. Larger networks exhibit a lower probability (at the same percentage of deletions) for the HD bump to get stuck (Supplementary Fig. 9).

Finally, the results from our model when it performed the behavioural task shed some light on the participants’ behavioural performance and the observed age differences. We found that young adults only had noise on the weights and background drive, while we added minimal neuron deletion for older adults. We replicated the delay effect, in which the angular error increased after the 20-second delay. This effect was larger when neuron deletion was involved, as in the participant data, where older adults performed worse. This is a mechanistically interpretable indication of neuronal death exacerbating the drift of the HD bump during the delay period, leading to deterioration in orientation coding, specifically in older adults. However, in the current simulation setup, we could not assess the effect of deteriorated HD in downstream areas known to carry HD information^8–12^. Future work should further explore these systems-level effects.

Several limitations of our study should be acknowledged. The precise architecture of the human HD system is not known, and certain compensatory mechanisms (e.g. homeostatic plasticity or recalibration) were not implemented. The model also does not simulate downstream structures that rely on HD signals, such as retrosplenial cortex or entorhinal cortex, where aging may have additional effects. Despite these limitations, the model reproduced key features of aging behaviour and offers testable predictions for future work, including broader HD tuning curves and increased drift in delayed-orientation tasks.

To conclude, we have presented a novel VR paradigm and behavioral evidence for deficits of HD coding in older adults, without response clustering. This data is consistent with a biologically plausible HD model that can accommodate aging-like perturbations and naturally accommodates vestibular drive. The latter carries HD activity across basins of attraction, consistent with our behavioural results. This 3-ring attractor model offers rich mechanistic insights, predicts broader tuning curves, and continuous drift in response to perturbations. Future work should investigate the detailed properties of the vestibular system as a driver of HD activity, as well as how age-related changes in the (sub)cortical areas that convey sensory information to anchor the HD signal may be selectively affected by aging.

## Methods

### Participants

Thirty healthy young adults aged between 18 and 30 years (23.77 ± 3.20, female n=17) and thirty healthy older adults aged between 60 and 80 years (68.31 ± 5.58, female = 15) participated in the study. Participants were fluent in German, did not report neurological or psychiatric disorders, and were not colourblind. The experiment was approved by the local ethics committee, and all research was performed in accordance with relevant guidelines and regulations. Written informed consent was obtained from each participant before starting the experiment. Participants were compensated financially for their participation. We excluded one older participant due to an incomplete dataset.

### Experimental design and procedure

Every participant came on two consecutive days (Fig. 1C). On the first day, participants performed three different colour tests (Ishihara test, colour arrangement, and colour matching) to verify that they were not colourblind and to assess their colour perception. Then, participants performed the learning task in immersive VR using a HMD, followed by the Heading Recall task (description in detail below). Each day, participants performed two blocks of the Heading Recall task with a break in the middle. After each block, they completed a task where they had to place the learned landmark. Participants returned on the second day to repeat the learning task and two new blocks of the Heading Recall task. Throughout the experiment, participants wore a HTC Vive HMD with a wireless adapter and could actively walk within a 5 by 5 metre space without constraint.

### Environment

The virtual environment was created using Unity (version: 2019.3.0f6). The ground had a grassy texture without any visible pattern, and the skybox represented a blue sky with no specific pattern. In addition, a cylindrical wall (diameter: 100 virtual meters (vm), height: 4.5 vm) surrounded the participant (Fig. 1A). The distance between the cylinder and the area where the participant could walk around was large enough, so the participant could not reach it. The texture applied to the cylinder was a colour gradient (Fig. 1A-B). Furthermore, a large black rectangular tower - acting as a distant landmark with a width of 34.8 vm - was placed outside the cylinder and centred at 0°. I.e., the landmark was always located at the same color value of the wall gradient. The landmark covered 10° of the participant’s field of view when they directly faced it from the centre of the environment. Its position was held constant across the whole experiment and between participants.

### Learning task

The goal of the *learning task* (Fig. 1D) was to learn the landmark’s position until meeting a predefined performance criterion. Participants had up to 100 trials to remember the correct position. On each trial, participants first walked to a random starting location indicated by a blue circle on the ground. Upon stepping on it, the cylindrical wall and the landmark disappeared. Next, a 3D arrow appeared in front of them, prompting the participants to rotate either left or right by a fixed amount: small (50°), medium (105°), or large (160°). After completing the rotation, participants heard an auditory cue and saw a text asking them to respond immediately. We added the auditory cue to ensure older participants would not overturn but stop at the correct orientation. To register their response, participants had to rotate physically until they faced the remembered position of the landmark. After rotating and confirming their orientation, participants received feedback in the form of smileys shown in front of them. Feedback was based on the absolute angular difference between the landmark’s actual position and the participant’s facing direction. They saw either a red (error >45°), orange (20°<error<45°), yellow (8°<error<20°), or green smiley (error<8°). To complete the learning, participants had to pass a threshold calculated by taking the average of the last five angular errors. If the average was below 8°, participants passed the learning phase and could proceed to the Heading Recall task. If not, they continued the learning task until they succeeded or reached 100 trials. Every participant passed the learning before the 100-trial threshold.

### Heading Recall task

The *Heading Recall task* (Fig. 1E) was conducted in the same environment, where only the cylindrical wall was visible, but not the landmark. However, note that during learning, participants may associate a color from the gradient with the landmark’s position. By doing so, participants using this strategy could treat it as a feedback signal during the Heading Recall task (when the landmark was not present). Participants started a trial by walking to a random location indicated again by a blue circle on the ground. Upon reaching it, a 3D arrow asked the participant to rotate until they faced the correct direction for the start of the trial. Next, they pressed the trigger button of the controller to start the first turn. The cylinder disappeared (the ground texture and the sky were still visible) when they pressed the trigger, and another 3D arrow instructed participants to rotate left or right by a predefined amount, similar to the learning task. The rotation was followed by a delay of 20 seconds for 50% of the trials. During this delay, participants were asked to perform a backward counting task with steps of 2, starting with a randomly generated number between 90 and 110. The goal was to prevent participants from rehearsing their answers during the delay period. The experimenter ensured that the participants were counting correctly during the whole task. After the delay or no delay, participants had to answer by physically rotating to face the landmark, similar to the learning task. However, participants received no feedback about their performance, the cylindrical wall reappeared, and the subsequent trial started.

The trial sequence was created prior to the start of the experiment and comprised 48 unique trials, each repeated three times. A unique trial consisted of a specific combination of rotation (small, medium, and large), delay condition (0 or 20 seconds), and the required rotation to face the landmark (0°, 45°, 90°, 135°, 180°, -135°, -90°, -45°). All 144 trials were separated into four blocks of 36 trials, with two blocks administered on each day. Finally, at the start of each block, participants were shown the correct position of the landmark for as long as they desired. The goal was to ensure participants could refresh their memory about the correct location of the tower before starting a new block.

### Tower placement task

At the end of each block, participants were asked to place the landmark within the environment. The goal was to obtain a readout of each participant’s memory of the landmark position and to ensure participants memorised the landmark position correctly throughout the experiment. During this task, they only saw the cylindrical wall and placed the landmark by facing the correct position and pressing the trigger button. Then, the landmark appeared at the indicated position, and participants could refine their answers by using the controller to move its position either left or right. Lastly, they locked their answer and continued with the Heading Recall task.

### Statistical analysis of behaviour

Statistical analyses - performed using Python (version: 3.7.4) and R (version: 4.1.3) - involved repeated-measures ANOVAs and t-tests. The learning task analysis was based on the number of trials completed by participants. The Heading Recall task was mainly based on absolute angular error, i.e., the difference between the landmark’s actual position and the participant’s facing direction when answering. For more details, see Supplementary Methods.

### The self-connected HD attractor model

To compare a biologically more plausible ring-attractor to classic HD circuit models, we first implement the latter. Our specific implementation follows ^30^. For the details of the architecture, see ^24^. Briefly: A single ring (Fig. 1H) with 361 neurons was implemented as a firing rate-based neuron model updated by angular velocity. The connection weights followed a Gaussian distribution for symmetric weights (recurrent bump-forming weights) and velocity-modulated asymmetric weights to shift the activity packet in response to angular velocity input during turns. This model employs mixed excitatory-inhibitory weights (violating Dale’s principle) and employs recurrent self-connections, requiring very precisely tuned connection weights to allow the attractor to self-sustain, but not saturate (activity spreading throughout the network) or collapse (due to lack of self-excitation). We customarily implemented visual feedback as a ring of visual neurons, each firing for a given view and providing topographic feedback to the HD ring.

### A robust multi-ring HD attractor model

The biologically plausible HD attractor architecture is a 3-ring structure, partly inspired by previous models^27,28,53^. Fig. 1F shows the architecture of the 3-ring model. Excitatory self-connections are removed, and instead, separate excitatory and inhibitory connections are used (conforming to Dale’s principle^54^). One excitatory ring projects to two inhibitory rings, which project back (CW and CCW, respectively) to the excitatory ring.

### The neural dynamics of the attractor network

To calculate the neural dynamics (in both HD models), we used a firing-rate neuron model. The activation of each neuron u within each ring is calculated according to the following differential equation:

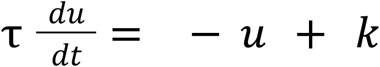

With

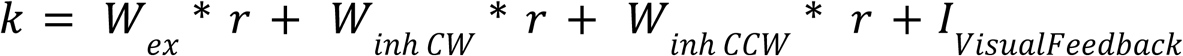

suitable discretized. *τ* is the neuronal time constant. *I* is the additional current used for the visual feedback (see below). *W* are the weight matrices, positive for excitatory or negative for inhibitory. Activation is mapped to the firing rate *r* with a sigmoid transfer function:

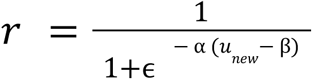

where α and β determine the sigmoid shape, this sigmoid normalises the firing rate to range from 0 to 1.

### Synaptic weights

Connection weights followed a Gaussian distribution, replicated for each neuron in each ring (Fig. 1G). For the excitatory ring projecting to the inhibitory rings, connections were topographic (e.g., neuron 100 maximally driving neuron 100 in both inhibitory rings, etc.). Inhibitory weights targeting the excitatory ring were offset to the right for the CW ring and to the left for the CCW ring. Together, the inhibitory connections from the CW and CCW rings to the excitatory ring neurons yield symmetric inhibition around the activity bump, leaving a zone of relative disinhibition (Fig. 1G). Since AHV cells^11,71^ were not explicitly modeled, the activity packet was translated by scaling inhibitory weights in a velocity-dependent manner, implementing a temporary asymmetry in the weights during turns, a proxy for velocity scaled firing rates^60^. This imbalance, left and right of the bump, corresponds to the direction in which inhibition is decreased. Changes to the multiplicative pre-factors of the inhibitory weights were fitted to obtain different translation speeds. The resulting speed profile of the bump of activity was computed to obtain the differences in inhibitory factors (*ΔF_inh_*) for changes in the angular head velocity of the agent (*V_ang_*):

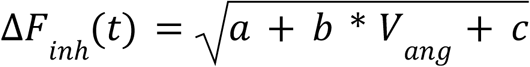

where parameters *a*, *b,* and *c* scaled the function to match the physical head velocity changes and the rotational speed of the bump.

Since there is no self-connection, the activity bump was supported and maintained by sending (constant) background drive to all three rings^27^, leading to the formation of the bump in the zone of disinhibition. The background drive followed a probability mass function similar to a Poisson distribution. This input can be thought of as the signal received from the vestibular system since lesions to the vestibular organs extinguish the HD signal^63,64^. Upon supplying moment-to-moment variation in the background drive, it leads to symmetry breaking, and the HD bump forms at a random location, mirrored on all three rings.

In addition, we modeled the visual input as described in previous studies^24,27,28,53^, where a separate “visual” ring had a one-to-one connection with the excitatory ring (Fig. 1F). Similar to previous models, these feedback connections stabilized the signal against drift (see below) by sending a small current to the neurons in the excitatory ring associated with the agent’s current heading.

To calculate the HD over the neuron ensemble, we used a similar equation to ^27^:

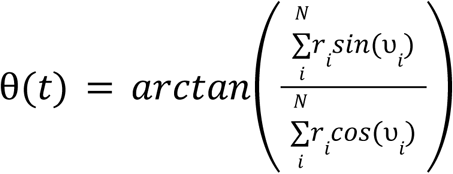

where *θ(t)* is the estimated population HD of the neuron ensemble *N* at time *t*, *r* is the rate value of the neuron *i*, and *ϑ* is its preferred heading direction.

### Sources of noise

We implemented different noise sources to characterize the robustness of both HD models. The first source was the amount of noise added to the Gaussian distributions of the weights - i.e., synaptic noise. The second source was associated with the background drive (not applicable to the self-connected model, as it does not require a background drive). Lastly, we implemented neuron deletion by randomly removing neurons. For the 3-ring model, this was done in all three rings. A deleted neuron would lose all its connections with other neurons and have a firing rate of 0 during a simulation.

### Modelling of the HD behavioural task

For the second part of this study, we implemented the task that participants performed in the model, applying several simplifications. Specifically, we simulated an agent that only performed turns but no translations, always located at the center of the environment. The colour gradient was mapped to the visual ring (Fig. 1F), providing feedback only when visible to the agent/participant. We hypothesized that the agent could (re)orient itself when looking at the colours, similar to how participants used the colours during the behavioral task. This was implemented by up-regulating the feedback connections between the visual ring and the HD ring when the participant could see the cylinder with the colours. Conversely, when the arena gradient disappeared, reversed feedback connections (from HD to the visual ring) were used to compute the colour the agent might estimate as coherent with the current facing-direction indicated by the HD activity. This implemented a readout of the agent’s answer. In addition, the agent was simulated to “believe” it was facing the landmark when the activity bump of the excitatory ring was within 356° to 4°, since every participant learned the position with an accuracy of 8°. We computed the agent’s physical heading (ground truth) and the estimate based on the HD neuron assembly at every step during a simulation.

We separated each trial into six different Phases (a-f) to replicate the trials participants performed in the behavioural task. After the last rotation from the agent at Phase f, the agent’s physical heading was recorded to measure the absolute angular error of the trial.

During Phases a, c, and c, the excitatory ring received visual (feedback) inputs since the arena with the colour gradient was present in the task (and not in Phases d, e, and f). However, the reversed connection profile from visual feedback to the HD ring allowed the agent to estimate the colour it was facing when the gradient was absent (see above), similar to the strategy reported by participants in the behavioural task.

Each agent had a trial sequence of 30 unique trials, with the following combinations of required rotations to answer: 0°, 40°, 90°, 135°, and 180°; delay: 0s and 20s; and first turn: 50°, 105°, and 160°. Every trial was administered twice, for a total of 60 trials. The order of the trials was shuffled across agents. Importantly, the model did not reset the rate of each ring between consecutive trials (i.e., the rate values at the end of a trial were carried over to the start of the subsequent trial), and the agent’s starting heading was the last one from the previous trial. This way, the visual feedback at the start of the trial had to reset the agent’s heading to the HD ring’s heading, as might be assumed for human participants.

In addition, to match the agent velocities to those produced by participants in the behavioral experiment, we created two distributions (Supplementary Fig. 8A-B): one for the velocity when participants performed the first turn (Phase d) and another for the velocity while answering (Phase f). Those distributions included both young and older participants. We removed the 5% most extreme values on each side of the distributions to eliminate outliers.

At the start of a trial, the model randomly picked a value from the two distributions. In addition, during the answer phase, participants could either rotate back toward the starting orientation (i.e., at Phase c) or continue in the same direction and complete the circle. We computed the percentage of circle completion for each participant (Supplementary Fig. 8C), and each agent followed probabilistically one of those percentages.

### Processing and Analysis of the Simulations

The analyses used Matlab (R2022a), Python (version: 3.7.4) and R (version: 4.1.3). The perturbations were implemented in the attractor models in 3 ways: deletions of cells (corresponding to neuronal loss or disconnection of a neuron from the attractor), noise in the incoming background drive, and static perturbations to the attractor weights on top of the Gaussian connectivity profiles. The analysis investigated the influence of the noise (single or combined) on static simulation and the model performing the HD behavioural task. For more details, see Supplementary Methods. In addition, details for the comparison between behavioral and modeling data (i.e., kernel density estimation for model-data fit) can also be found in the Supplementary Methods.

## Supporting information

Supplementary Informations

## Acknowledgements

We gratefully acknowledge M. Schaumburg, C. Winter, and F. Zeller for their help in the data collection. We also thank A. Imtiaz for his advice on creating the behavioural task. T. Wolbers and M. Bernard were supported by the Deutsche Forschungsgemeinschaft (DFG, Project-ID: 425899996 – SFB 1436) and by the European Social Funds (ESF) (Sachsen-Anhalt Wissenschaft Spitzenforschung/Synergien: HD-CODING). A. Bicanski acknowledges funding from the Max-Planck Society. Last, we thank the participants for making this work possible.

## Contributions

Matthieu Bernard (MB) completed this project in collaboration with Jonathan Shine (JS), Andrej Bicanski (AB), and Thomas Wolbers (TW). MB, JS, AB, and TW conceptualised the work. MB managed project administration, programmed the VR task, and collected behavioural data. MB and AB developed the computational head direction model. MB analysed the data and visualised the results. AB and TW jointly supervised the work. MB drafted the manuscript. MB, JS, AB, and TW revised it.

## Supplementary information

See supplementary document.

## Notes

### Competing Interest Statement

The authors have declared no competing interest.

