## Supplementary Informations for "How human aging disrupts the head direction network: evidence from VR experiments and mechanistic models"

### Supplementary Methods

#### Statistical analysis of behaviour

For the learning task, we extracted the number of trials completed by each participant, separately for each testing day. We also extracted the landmark placement for each participant.

For the Heading Recall task, we calculated the absolute angular error similarly to the learning task, i.e., the difference between the landmark's actual position and the participant's facing direction when answering. However, we corrected those values by taking the position as indicated by the participant in the tower placement task and using it as the reference when calculating the angular error. In addition, we recorded the heading of the participant during the delay period and calculated its circular standard deviation. The goal was to assess if participants moved their heads during the delay period.

Further analyses addressed the angular error on trials where participants had already faced the landmark before answering. Since the correct answer for those trials was to respond without rotating, the analysis focused on the impact of delay and the first turn. Hence, we quantified the rotation performed by the participant. For each trial, we checked if the movement was below 5° and calculated the percentage of time when the participant moved. We decided to set 5° as the threshold because participants typically make slight head movements even when not asked to rotate, as seen during the delay period. Lastly, to investigate if there was a clustering of answers, we extracted the final heading of the participant after answering, and calculated the circular standard deviation across the different repetitions of the unique trials.

Finally, we calculated the scores for each control colour test for each participant. For the Ishihara test, we computed the percentage of successful trials. For the test assessing colour arrangement (i.e., the placement of colours to form a gradient, such as red to green), we computed the number of trials required to obtain a perfect answer for each arrangement.

Then, we averaged those values to create a percentage of accuracy. For the colour-matching tests, we calculated the mean across trials for each difficulty (easy, medium, and hard). Next, we either averaged those values to create an overall colour-matching score or separated them to get a score for each difficulty.

#### Tracking experimentally recorded HD with the model.

In order to map the behavioural results and performance of human participants onto the mechanistic model, we tested whether the HD of the excitatory bump followed changes in velocity and direction of participants from the behavioural task by extracting the movement trajectories over two minutes. Our model followed trajectories without drift when visual feedback was present (Fig. 3B). In accordance with past studies, when removing the visual feedback, drift accumulated between the real HD and the HD of the attractor.

#### A recurrent HD attractor cannot accommodate age-related changes.

To establish a baseline, we subjected the standard model of HD (one ring with recurrent weights) to the type of noise outlined above: deviations from theoretically perfect synaptic weights, deletions (neuron deletion and/or disconnection), and noise in the network inputs. In his seminal study, Zhang outlined a possible effect of noise on the weights but did not assess the impact of deletions[1]. We replicated this recurrent 1-ring model as implemented in [2], removed a single neuron from the attractor (red line in Fig. 3A), and let the bump move in one direction. We found that when the bump reached the dead neuron, the bump would collapse, and the signal would flatten (Fig. 3A). In addition, we performed the same simulation with the 3-ring model while having 1% of neuron deletion (red lines in Fig. 3A). The simulation showed that the bump completed a full circle by passing through the dead neurons without getting flat (Fig. 3A). This result provided evidence that a self-connected attractor can stop functioning with a minor disruption. We confirm the model's original assessment of limited noise tolerance in this model architecture, which is dependent on recurrent connections. Finally, variations in inputs cannot be studied in the self-connected attractor as activity is self-sustained.

### Processing and Analysis of the Simulations

#### A / *Assessing drift, reduction of representational space, and tuning curve width*

We implemented perturbations to the attractor models in 3 ways: deletions of cells (corresponding to neuronal loss or disconnection of a neuron from the attractor), noise in the incoming background drive, and static perturbations to the attractor weights on top of the Gaussian connectivity profiles.

To assess the impact of noise on the model, we measured HD cell peak firing rates and their variance, the full-width-half-maximum (FWHM) of the tuning curves, and the drift between the real HD and model HD. We compared these measures for different types/levels of noise. Broad tuning curves give a more imprecise heading signal.

#### B / Reproduction of the behavioural task using the model and analysis of its simulations

Testing the simulated behavioral task revealed a higher propensity for a perturbed attractor to occasionally get stuck at smaller network sizes. That is, a bump can be present, but it fails to react to velocity inputs. Thus, we conducted a separate analysis to quantify the percentage of time the network would get stuck and the influence of angular velocity inputs. In this analysis, we ran the behavioural task using the same first turn (105°). One hundred agents with damaged ring attractors repeated each condition three times. The conditions varied the number of neurons in each ring (361 or 721) and the percentage of deletions (1% to 15% for 361 neurons and 1% to 30% for 721 neurons). The head turn velocity when the agent answered (Phase f) was also varied across conditions: it was either set as the average value from the distribution or the average value with the addition/subtraction of one/two standard deviations. For each trial, the simulation ran until an answer was given by the agent or until the activity bump got stuck in a local minimum while answering. The probability of trials getting stuck in local minima across conditions was computed after all simulations were run. From the results of this analysis (see supplementary Results section), we decided to use a network with 721 neurons, which approximates the order of magnitude of HD neural on the rodent attractor[3] and reduces potential confounds due to numerical limits in small networks.

To investigate the impact of different noise sources on the performance during the behavioural task, we systematically varied the amplitude of each noise variable, either individually (Supplementary Table 3) or combined (Supplementary Table 4). For each noise level, 30 agents performed the task and used a modified trial sequence. These simulations were characterized by the following metrics across the different phases of the trials: *angular error* (difference between the agent's HD and the HD indicated by the network at the end of a phase) and the *rate of change* of the HD (average of the difference in velocity of the agent's HD and the bump). The final rate of change values would either be positive if the agent's velocity was faster or negative when the velocity of the bump was faster on average.

To compare the human data and agent data simulated with the behavioural task, we used the average angular error measured at the end of the trial for the different expected rotations to the landmark (see Statistical analysis of behaviour). The range of error was different between human and agent data. However, despite the relative leap in biological plausibility, the model is missing many aspects of biological brains, and tuning parameters to match the absolute numbers in the behavioral data is of limited utility. E.g., the present model omits the extensive (sub)cortical HD circuitry (RSC, Subicular cortex, thalamus) that is involved in conveying feedback back to the attractor (see [2,4–6]), and could be independently affected by aging. Instead, to compare the model and the data, we mean-scaled the behavioral data and model-data separately for each age population and delay condition. Next, we subtracted each average for the different expected rotations from the landmark between the human data and the agent data (for each delay condition and age group separately). Lastly, we took the absolute values of the data and averaged them to obtain a match value between the profiles. This analysis was done by including all expected rotations for the landmark and excluding the 0 ° rotation.

#### C / Analysis to match behavioural and modelling data

To find the best match between the behavioural data and the model, we run several simulations with combinations of noise types (Supplementary Table 4). To select a model configuration, we treat young adults with a 0-second delay as the baseline and select the best model according to the criteria described below. To choose the "aged model", we refer to the increase in error observed between young and old human subjects. Thus, the model selected as a candidate for the aged condition is based on the relative error increase in humans (approximately 50%), added to the ground-truth model candidate. In addition, we excluded the 0° rotation to find the best match, since slow velocities are more prone to error. We account for the relative change between young and older adults and between delay and no-delay conditions by rescaling the simulated data to the same mean (mean scaling) to compare the shapes of the bar plots across age groups and delay conditions separately. We refer to the difference in profile shape between the behavioural and model data as dShape, and to the relative increase in error as dError. Next, these two measures can be weighted to obtain a compound score: MatchScore = Alpha * dShape + Beta * dError; with Alpha = Beta

= 0.5 to give equal weight to both scores. The MatchScore obtained is the basis for the descriptive analysis.

In addition, we calculated the model-data fit using log-likelihood using kernel density estimation. First, we separated the young and older adults' data to find their best matches separately. For each model (combination of noise levels, see Supplementary Table 4), we employed kernel density estimation applied to the absolute angular error of the simulated data for each trial type (combination of delay condition and expected rotation to the landmark). This yields an empirical error distribution for each simulated model under which we assess the log likelihood of the real data. Then, we interpolated the real data from this distribution and calculated its logarithm for each trial type. Therefore, each real data gives a likelihood that was summed for each trial type. This gives the total log-likelihood of the real data under a particular simulated model, for both young and older adults' data. Moreover, we compared several bandwidth heuristics: Unbiased cross-validation, Biased cross-validation, and Sheather-Jones, and they all produced the same results. For the results, we showed the results for the unbiased cross-validation.

### Supplementary Results

#### Performance in color perception tests was comparable between age groups.

To ensure that differences in colour perception did not confound our results, we first compared the performance in the colour tests between the two age groups using Welch's t-tests. For the Ishihara test, we found no significant difference ($t(54.99)=0.23$, $p=0.81$) between older (mean=98.38 ± 3.60) and younger (mean=98.17 ± 3.14) adults. In contrast, younger adults (mean=98.96 ± 1.14) performed slightly better than older adults (mean=96.30 ± 4.17) in the arrangement test ($t(32.13)=-3.31$, $p=0.002$). Note, however, that both groups performed at very high levels (i.e., > 95%). Lastly, a significant difference was found in the colour-matching test when all difficulty levels were included ($t(40.77)=3.25$, $p = 0.002$). However, the colour gradient mapped onto the circular arena was bright and more closely related to the task's easy difficulty level. We therefore ran another t-test including only the easy difficulty, which yielded no difference ($t(54.04)=-0.87$, $p = 0.39$) between older (mean = 32.48 ± 9.91) and young participants (mean = 29.96 ± 12.01).

#### No performance differences across testing days

The experiment was performed across two consecutive days. To verify that data from both sessions could be pooled, we compared the mean corrected angular error for Day 1 and Day 2 (Supplementary Fig. 1). A two-way ANOVA showed a main effect of age ($F(1,57)=9.83$, $p=0.003$) but no main effect of days ($F(1,57)=2.11$, $p=0.15$) nor interaction ($F(1,57)=0.0002$, $p=0.99$). This result suggests that task performance was stable across the testing days, enabling us to perform subsequent statistical analyses on the combined dataset.

#### The 20-second delay impaired performance in both groups when facing the correct orientation before answering

We asked whether participants tended to remain stationary on trials in which they were already facing the landmark (Fig. 2F). In these trials, the optimal strategy is to hold the initial heading throughout the response interval— we reasoned that any additional movement might indicate orientation drift during the delay. For each participant, we therefore computed the proportion of trials on which the total rotation during the response phase was < 5° (horizontal black line in Fig. 2F). We selected a 5° threshold, rather than a strict 0°, to accommodate inadvertent micro-movements that might escape participants' awareness. This was assessed by testing whether participants moved their heads during the delay phase (Supplementary Fig. 4). The average range of motion did not differ between age groups (young: 2.03° ± 5.26°; older: 1.67° ± 2.97°; $t(53.27) = 1.42$, $p = 0.16$).

After grouping the proportion of "fixed-heading" trials by the magnitude of the initial turn (Supplementary Fig. 5), a three-way ANOVA revealed significant main effects of age ($F(1, 57)=4.18$, $p = 0.045$), delay ($F(1, 57)=10.54$, $p = 0.002$), and first-turn size ($F(2, 114)=11.08$, $p < 0.001$). Significant interactions were observed for age × delay ($F(1, 57)=4.34$, $p = 0.042$) and delay × first-turn size ($F(2, 114)=5.54$, $p = 0.005$), whereas the age × first-turn interaction was not significant. Collapsing across first-turn categories showed that the 20-s delay reduced the proportion of fixed-heading trials in older adults ($t(28)=2.93$, $p = 0.007$) but not in younger adults ($t(29)=1.32$, $p = 0.20$). Without a delay, younger adults exhibited a lower proportion of fixed-heading trials than older adults ($t(44.86)=–2.57$, $p = 0.013$); this age difference disappeared after the 20-s delay ($t(48.20)=0.61$, $p = 0.55$). Together, these findings indicate that the delay period caused greater orientation drift in older than in younger participants. However, this effect may be partly explained by the task design, which instructed participants to physically rotate on each trial, potentially reducing the likelihood of true 0° responses, as participants could expect to need to rotate on every trial.

#### Neuron deletions, velocity strength and size of network shape local attractor basins.

The static simulation results (cf. Fig. 4) suggested that the bump moved and eventually reached a stable position as neuron deletion increased. That is, deletions lead to the emergence of attractor basins, and initial HD settles into those local energy minima of the network. However, those simulations were performed without velocity inputs. Since we aimed to subject the attractor to the behavioral task performed by the participants (trials similar to those of actual participants), we tested the behavior of the perturbed/damaged attractor with a moving packet of activity. We quantified performance as the probability that the bump would settle into a local minimum (i.e., remain stationary for at least 1 second while a velocity input was present, i.e., during the answer phase).

This probability of settling in local minima increased with the number of deletions (Supplementary Fig. 9), and was attenuated when the total number of neurons in the attractor was expanded, while keeping the percentage of inactive neurons constant (Supplementary Fig. 9). By doubling the number of neurons in the network (from 361 to 721), we obtained the same distribution and range of values when the percentage of deletions doubled (from 15% for 361 to 30% for 721; Supplementary Fig. 9). That is, a large network can tolerate the same percentage of noise better than a small network, because there are more combinatorial possibilities to distribute deletions among a larger number of cells. This has direct implications for the design of biologically plausible attractor networks across species (see Discussion).

Finally, the probability of settling in local minima depends on the velocity input received. The mean velocity value across the rotations during the response phase varied. For example, this value was larger for 180° rotations than for 45° rotations (see Supplementary Fig. 8). We observed an inverse bell-shaped distribution, with the probability of getting stuck lower for 180° rotation and higher for 45° rotation (Supplementary Fig. 9). In addition, we varied the rotation velocity and found that, at slower velocities, the probability of getting stuck in local minima was higher than at faster velocities (Supplementary Fig. 9). This observation held for different percentages of neuron deletion (Supplementary Fig. 9). This suggests that a higher turning velocity in the attractor can successfully carry the activity packet across local minima.

#### Descriptive analysis of model-data fit

The process is based on several hypotheses and the analysis of our data so far: 1) The aging literature suggests there is little loss of neurons during aging, suggesting that our "aged model" should at most have a low percentage of deletions. 2) We did not find any clustering in our behavioural data, suggesting the model should not get stuck in local minima when velocity is present, which necessitates moderate to high variability in the background drive (to carry the HD bump over local minima). From the analysis of individual noise types, we find that weight noise and neuron deletion create attractor basins. 3) Young adults should have some background noise (needed for the plausible attractor to function) and low to moderate synaptic noise (accounting for the idea that weights cannot be 100% precise). 4) In comparison, a model simulating an aging agent should be at least as perturbed (in terms of weights and background drive) as the young model, and exhibit low neuron loss.

When running the simulation, we found that the range between the behavioural and model data increased with higher levels of noise. However, how the model fares in accounting for relative changes between young and older adults and between delay and no-delay conditions can be assessed. Therefore, we decided to rescale the simulated data to the same mean (mean scaling) to compare the shapes of the bar plots separately for age groups and delay conditions. The selection of the best fit was then based on both measures: relative error increase and profile shape, resulting in a match score with a lower value indicating a better match.

To select a model configuration, we treat the young adults' data with no delay as the baseline and select the best model. To select the “aged model”, we refer to the increase in error observed between young and old human subjects. Thus, the model selected as a candidate for the aged condition is drawn based on the relative error increase in humans (approximately 50%) added to the ground truth model candidate, while respecting conditions above. Once a candidate young adult's no-delay model is selected, it is applied to the young adults for the 20s delay, and the choice of older adult models is dictated by the closest match to the observed 50% increase in error for older vs. young. Finally, one last important value to measure is the scale factor between the age groups and the delay conditions. This way, we could check whether the overall range of values between the young and the older is close. Same for the delay condition.

First, we aimed to obtain a baseline by finding the best match for the young adults in the no-delay condition. We found that the best match score (3.26) across the simulation was for 10% WN + 10% BG. Next, we assessed whether the same model also provided the best fit for the 20s delay condition. When using the same approach, we found that the model with 10% WN + 20% BG had the best match score (5.70), while 10% WN + 10% BG had the 2nd-best (6.35). To decide the best fit, multiple approaches were taken to be as exhaustive as possible: smallest scores for 0s (3.26) and 20s (5.70); sum of the scores between 0s (9.62) and 20s (9.43); scale factor between delay conditions for each fit 10% WN + 10% BG (1.42), 10% WN + 20% BG (1.2). Among these approaches, the model fit for the 20s was the closest for 2/3 of the measures. Therefore, the model fit chosen for young adults is 10% WN + 20% BG.

Then we move on to model selection for older adults. Based on the hypothesis above, we looked only at noise levels equal to or higher than the one we found for young adults. We used the same methods, and for 0s the best fit would be 10%WN + 20%BG + 1%ND with a match score of 10.14. Finally, we looked at the 20s delay and found a best-fitting model that differed from 0s. The best fit for 20s was for 10%WN + 20%BG + 2%ND with a match score of 21.32. In contrast, the 2nd best for 10%WN + 20%BG + 1%ND has a match score of 22.28. To decide the best match, we applied the same approach as above: the smallest score for 0s (10.14) and 20s (21.32); and the sum of scores for 0s (32.82) and 20s (37.33). Scale factor for 10%WN + 20%BG + 1%ND (1.71) and 10%WN + 20%BG + 2%ND (1.22). In that case, the model fit for 0s was the closest for 2/3 of the measures. Hence, the model fit chosen for older adults is 10% WN + 20% BG + 1% ND. Therefore, we found the best possible match between behavioural data and model simulation. Interestingly, the main difference was the addition of minor deletions of neurons.

#### Log-likelihood analysis of model-data fit

To complement the hypothesis-driven choice of model based on descriptive statistics, we also calculated the log-likelihood of the real data given the error distributions of the models. After running the log-likelihood analysis, we found that the best-fitting model for young adults was 10% WN + 20% BG, and for older adults, 5% WN + 20% BG + 1% ND. While the best-fitting model for young adults is the same as in the descriptive analysis, the weight noise is slightly different for older adults. Thus, both methods yield similar results, albeit not strictly identical. However, across the board, log-likelihoods are lower in models without neuron death compared to those with neuron death, indicating that the ability of the neural model to accommodate neuron death is indeed crucial.

### Supplementary Tables

| Parameters | Values for 361 network | Values for 721 network |
| --- | --- | --- |
| Number of neurons on each ring | 361 | 721 |
| τ | 0.04 | 0.04 |
| dt | 0.002 | 0.002 |
| α | 0.7 | 0.7 |
| β | 6 | 6 |
| a | 146.20 | 1.36 |
| b | 47.70 | 2.49 |
| c | -12.09 | -1.17 |
| Factor Excitatory | 0.45 | 0.225 |
| Factor Inhibitory CW | 1.15 | 0.575 |
| Factor Inhibitory CCW | 1.15 | 0.575 |
| Factor Feedback | 5 | 5 |
| Background drive Excitatory | 28 | 28 |
| Background drive Inhibitory | 0.15 | 0.15 |

**Supplementary Table 1: Model parameters for the 361 and 721 neuron networks.**

| Weight Noise | Background Noise | Neuron Deletion |
|---|---|---|
| 0 | 0 | 0 |
| 5 | 10 | 5 |
| 10 | 20 | 10 |
| 15 | 30 | 15 |
| 20 | 40 | 20 |
| 25 | 50 | 25 |
| 30 | 60 | 30 |

**Supplementary Table 2: Percentage of noise for individual static simulations without velocity inputs.**
Yellow cells, simulation ran only for analysis of the shape of the tuning curve.

| Weight Noise | Background Noise | Neuron Deletion |
|---|---|---|
| 0 | 0 | 0 |
| 2.5 | 5 | 1 |
| 5 | 10 | 3 |
| 7.5 | 15 | 5 |
| 9 | 20 | 7 |
| 12 | 25 | 10 |
| 15 | 50 | 13 |

**Supplementary Table 3: Percentage of noise for individual task simulation.**

| **Weight Noise** | **Background Noise** | **Neuron Deletion** |
|---|---|---|
| 5 | 10 | 0 |
| 5 | 10 | 1 |
| 5 | 10 | 2 |
| 5 | 20 | 0 |
| 5 | 20 | 1 |
| 5 | 20 | 2 |
| 10 | 10 | 0 |
| 10 | 10 | 1 |
| 10 | 10 | 2 |
| 10 | 20 | 0 |
| 10 | 20 | 1 |
| 10 | 20 | 2 |
| 0 | 10 | 1 |
| 0 | 10 | 2 |
| 0 | 20 | 1 |
| 0 | 20 | 2 |

**Supplementary Table 4: Percentage of noise for combined task simulation.**

| Weight Noise | Background Noise | Neuron Deletion | Young adults | Older adults |
|---|---|---|---|---|
| 5 | 10 | 0 | -22571.99 | -29293.57 |
| 5 | 10 | 1 | -19132.82 | -19459.01 |
| 5 | 10 | 2 | -19713.80 | -19597.61 |
| 5 | 20 | 0 | -21594.84 | -27701.75 |
| 5 | 20 | 1 | -19335.44 | **-19437.84** |
| 5 | 20 | 2 | -20739.91 | -20391.07 |
| 10 | 10 | 0 | **-18692.85** | -21429.11 |
| 10 | 10 | 1 | -20015.79 | -19978.37 |
| 10 | 10 | 2 | -20767.87 | -20279.11 |
| 10 | 20 | 0 | -18493.65 | -21016.45 |
| 10 | 20 | 1 | -19877.72 | -19935.52 |
| 10 | 20 | 2 | -20624.01 | -20198.43 |
| 0 | 10 | 1 | -18728.48 | -20006.76 |
| 0 | 10 | 2 | -20558.34 | -20193.50 |
| 0 | 20 | 1 | -19011.66 | -19641.68 |
| 0 | 20 | 2 | -19688.12 | -19711.99 |

**Supplementary Table 5: Log-likelihood results.**

### Supplementary Figures

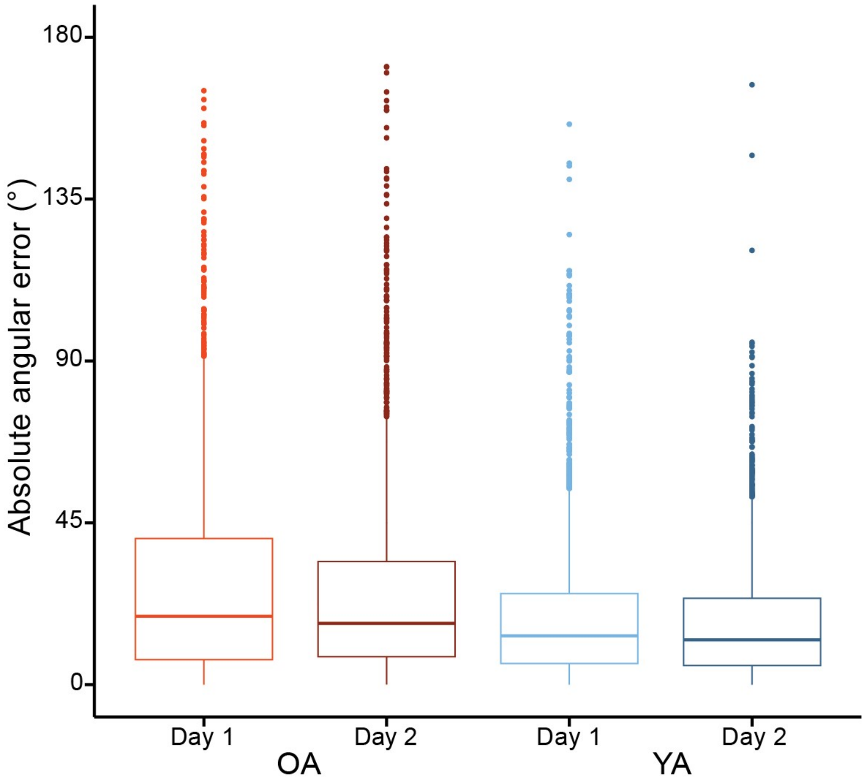


**Supplementary Figure 1: Control measures for equal performance between testing days**

Corrected absolute angular errors separated between age groups and testing days. OA, older adults; YA, young adults.

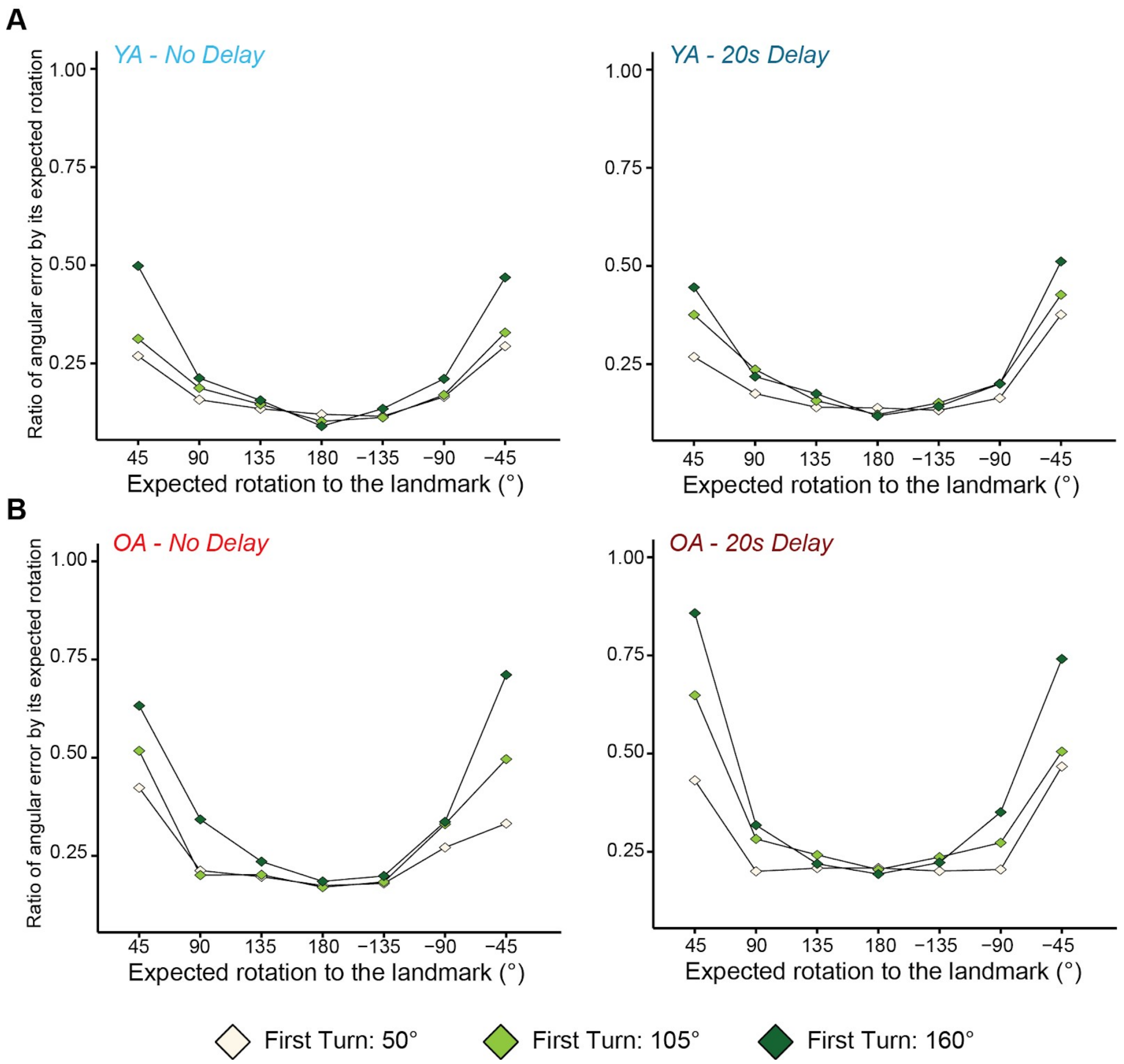


**Supplementary Figure 2: Ratio of angular error by expected rotation across delay conditions and age groups.**

Average ratio of the absolute angular errors divided by the associated expected rotation to the landmark for young and older adults for the two delay conditions. The ratio is separated by the first turn executed (coloured dots) for young (B) and older (C) adults after the delay conditions.

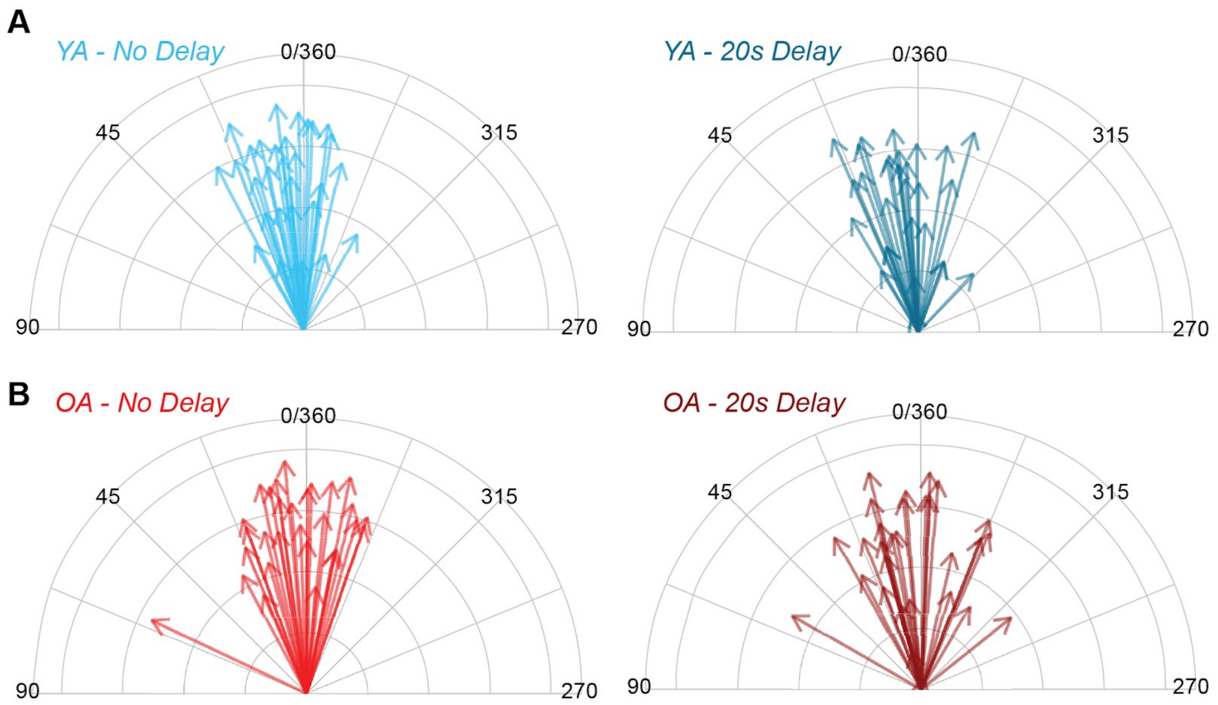


**Supplementary Figure 3: Answer distribution of participants when already facing the landmark before answering.**

Each arrow represents the average facing answer for (A) young and (B) older participants for (left) no delay and (right) 20s delay conditions. The length of the arrow represents the inverse circular standard deviation, with a shorter length describing a larger standard deviation.

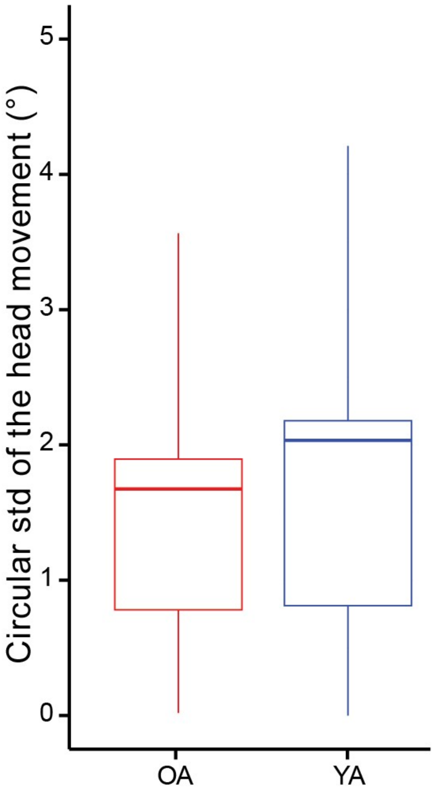


**Supplementary Figure 4: Control measures of head movement during the delay phase.** The circular standard deviation of the head movement performed by the participants during the delay period. OA, older adults; YA, young adults.

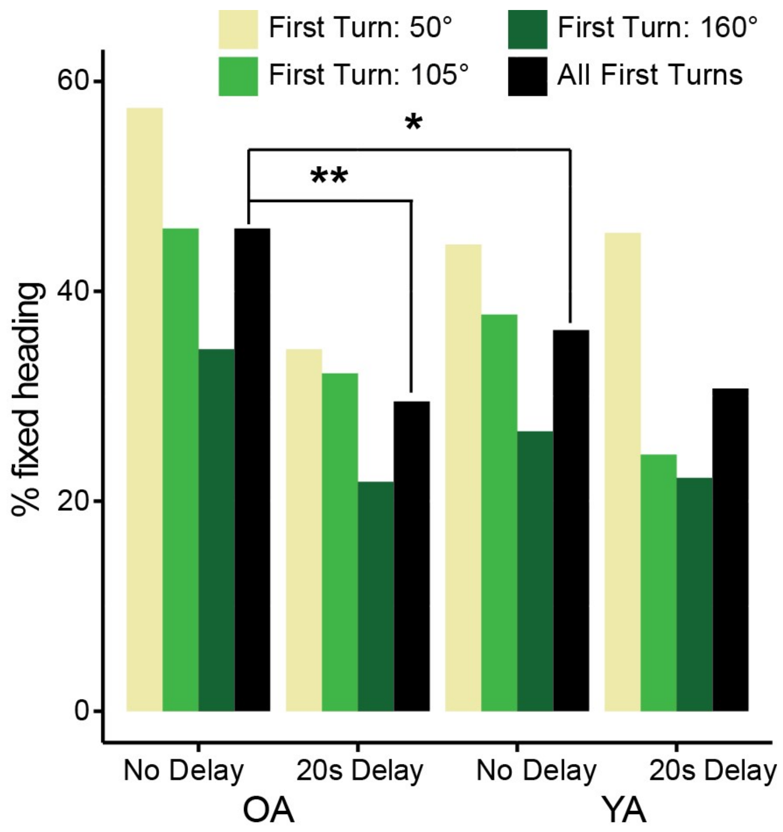


**Supplementary Figure 5: Fixed heading across age groups and delay conditions**
Bars represent the percentage of trials where participants did not move across age groups and delay conditions. Data were separated by the first turn performed before the delay condition. OA, older adults; YA, young adults.

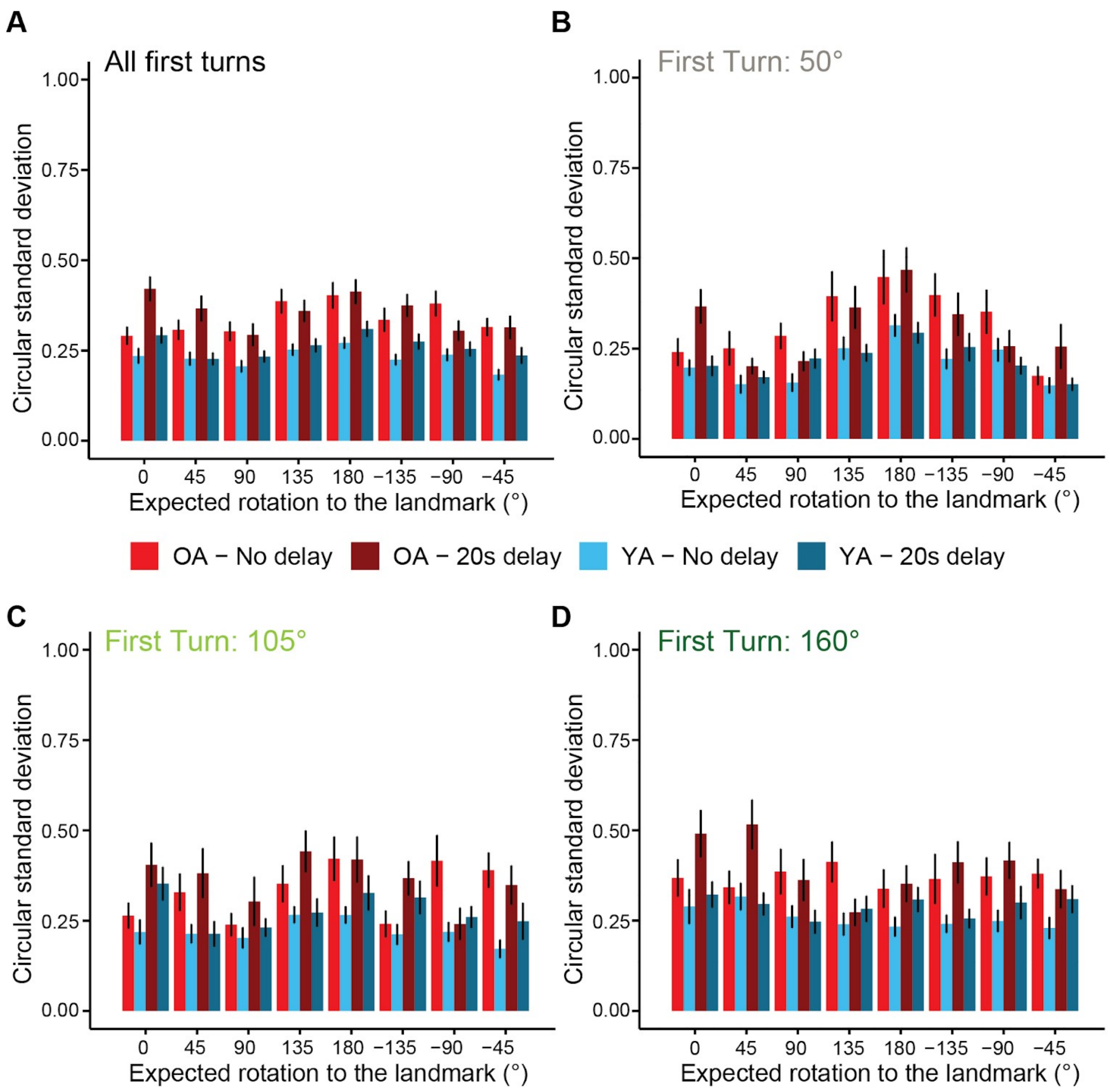


**Supplementary Figure 6: Checking the reduction of the representational space hypothesis.**

To check for potential clustering of answers, the circular standard deviation was calculated for young (YA) and older (OA) adults for the two delay conditions. The data were either (A) combined or separated by the first turn (B, C, D) performed. The bars represent the average circular standard deviation. The black line represents the standard error of the distribution.

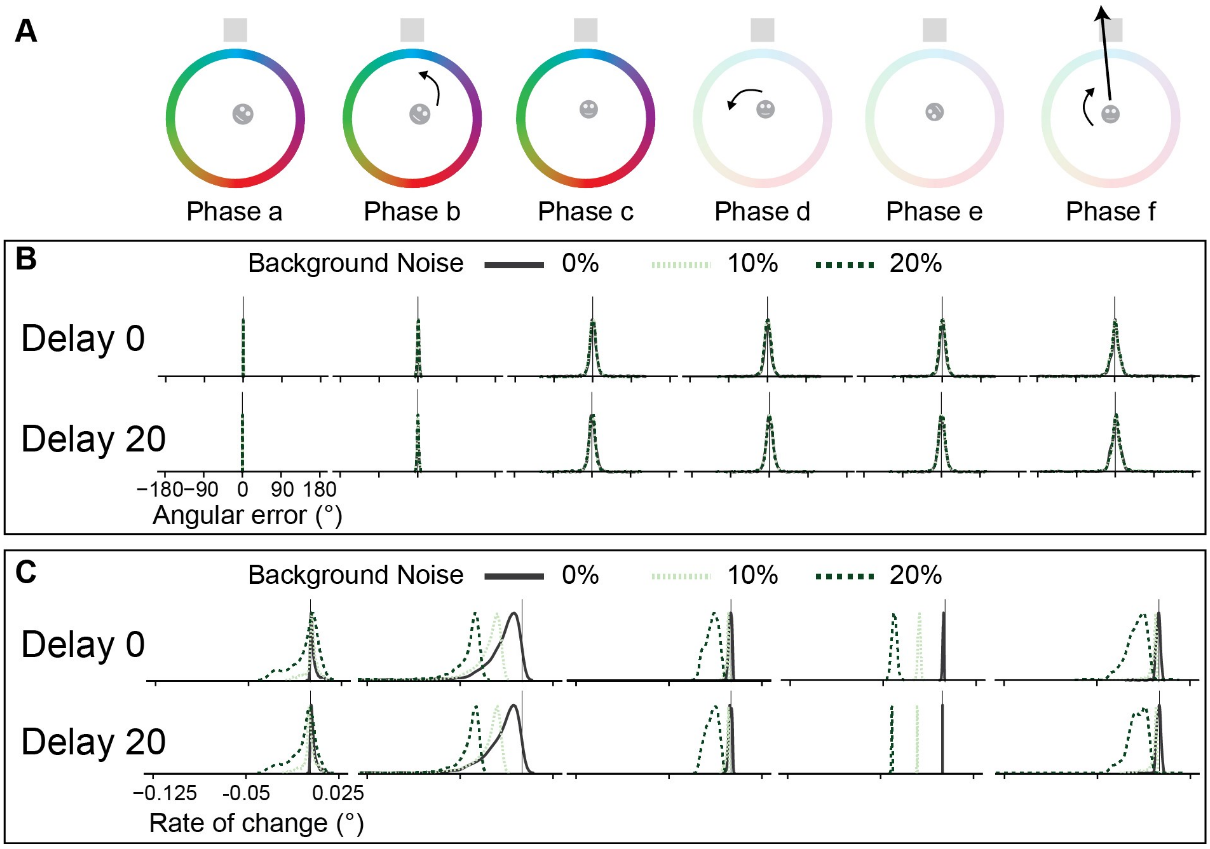


**Supplementary Figure 7: Evolution of angular error and rate of change for increased background noise.**

A, the different phases forming a single trial executed by the agent. Phase a, reset of the network; Phase b, rotation toward starting orientation; Phase c, facing starting orientation; Phase d, performing the first turn; Phase e; delay condition; Phase f; answer from the agent. B, the density plots show the distributions of the angular error (difference between agent HD and the HD indicated by the network at the end of a phase) measured at the end of the trial phases for background noise. The vertical black line represents 0. The plots were separated by level of noise (colours), by phases (from left to right), by delay conditions (top - no delay, bottom - 20s delay). C, the density plots show the distributions of the rate of change (average of the difference in velocity of the HD of the agent and the bump) measured for background noise. The vertical black line represents 0. The plots were separated by level of noise (colours), by phases (from left to right), by delay conditions (top - no delay, bottom - 20s delay).

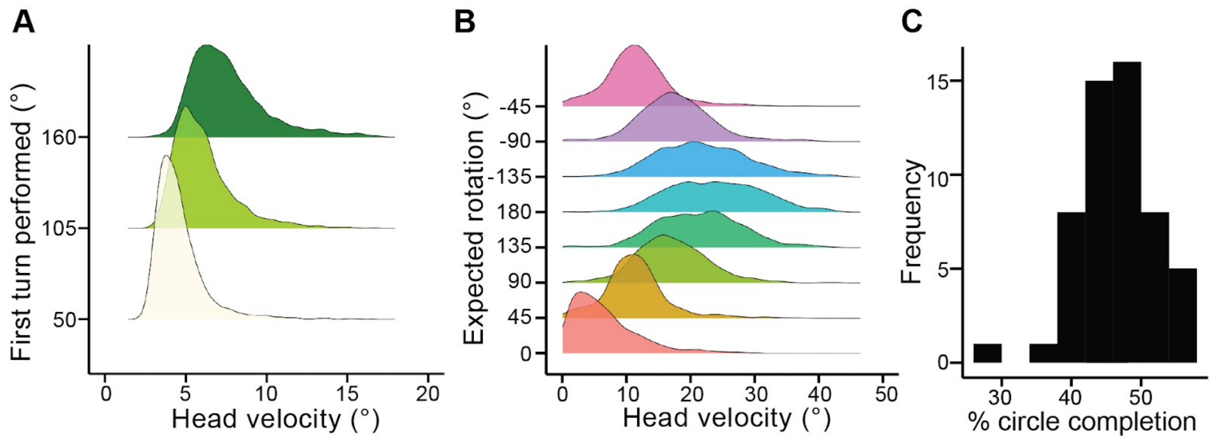


**Supplementary Figure 8: Participant distributions used in the model.**
Head velocity distributions of participants are separated by (A) the first turn performed and (B) the expected rotation to the landmark. C, the percentage of time participants performed a full circle while answering.

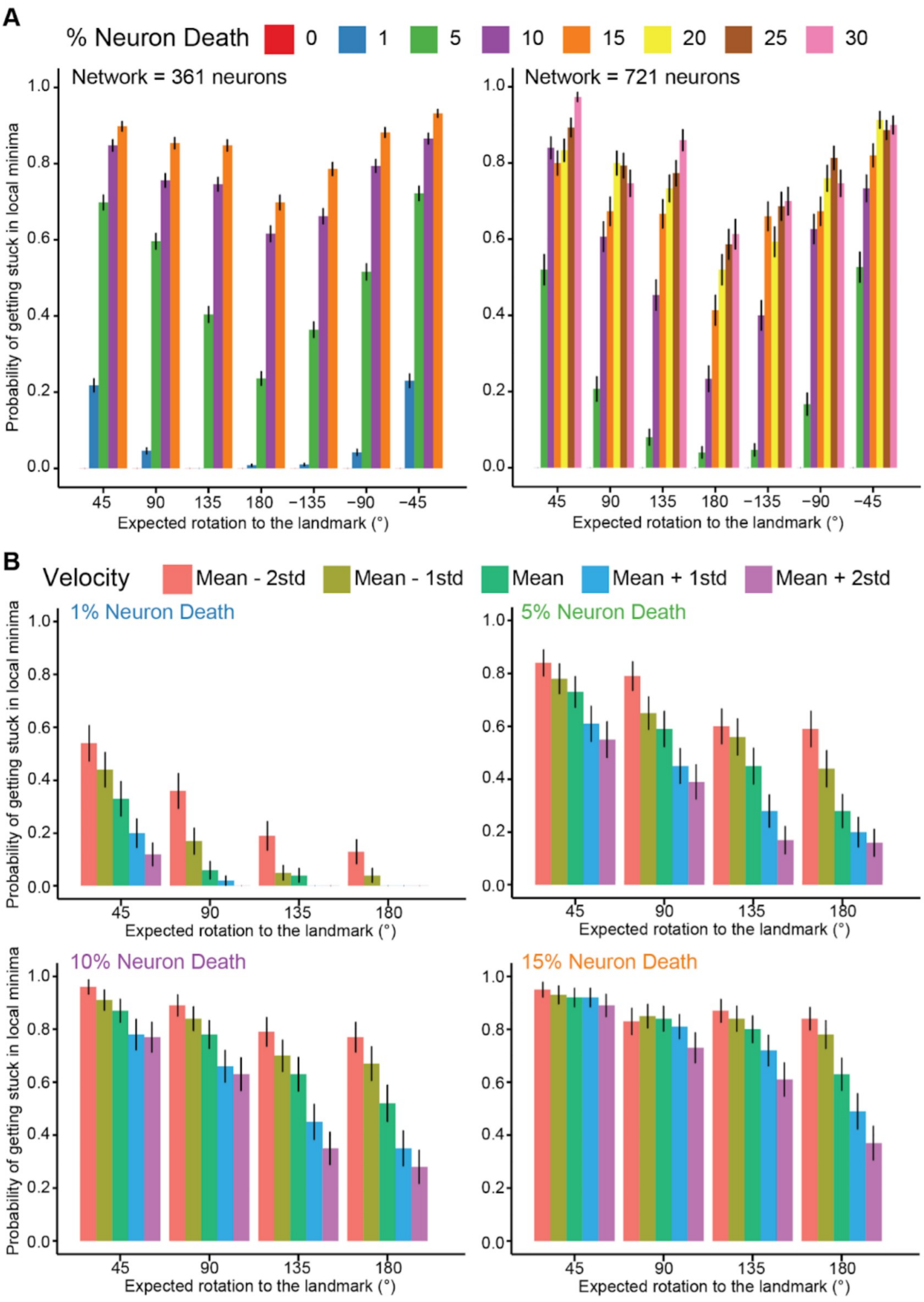


**Supplementary Figure 9: Probability for the attractor to get stuck in local minima when performing the task.**

A, average probability for the attractor with (left) 361 and (right) 721 neurons separated by expected rotation to the landmark. The maximal percentage of neuron deletion was set to 15% for 361, and 30% for 721 neurons. B, average probability for the attractor with 361 neurons separated by expected rotation to the landmark. The different colours represent the velocity used during the answer phase. Neuron deletion was set to (top left) 1%, (top right) 5%, (bottom left) 10% and (bottom right) 15%. The black lines represent the standard error.
